# Chalkophore mediated respiratory oxidase flexibility controls *M. tuberculosis* virulence

**DOI:** 10.1101/2024.04.12.589290

**Authors:** John A. Buglino, Yaprak Ozakman, Chad E. Hatch, Anna Benjamin, Derek S. Tan, Michael S. Glickman

## Abstract

Oxidative phosphorylation has emerged as a critical therapeutic vulnerability of *M. tuberculosis* (*Mtb*). However, it is unknown how intracellular bacterial pathogens such as *Mtb* maintain respiration during infection despite the chemical effectors of host immunity. *Mtb* synthesizes diisonitrile lipopeptides that tightly chelate copper, but the role of these chalkophores in host-pathogen interactions is also unknown. We demonstrate that *M. tuberculosis* chalkophores maintain the function of the heme-copper *bcc:aa_3_* respiratory supercomplex under copper limitation. Chalkophore deficiency impairs *Mtb* survival, respiration to oxygen, and ATP production under copper deprivation in culture, effects that are exacerbated by loss of the heme dependent Cytochrome BD respiratory oxidase. Our genetic analyses indicate that maintenance of respiration is the only cellular target of chalkophore mediated copper acquisition. *M. tuberculosis* lacking chalkophore biosynthesis is attenuated in mice, a phenotype that is also severely exacerbated by loss of the CytBD respiratory oxidase. We find that the host immune pressure that attenuates chalkophore deficient *Mtb* is independent of adaptive immunity and neutrophils. These data demonstrate that chalkophores counter host inflicted copper deprivation and highlight a multilayered system by which *M. tuberculosis* maintains respiration during infection.

## Introduction

Bacterial pathogens are subjected to diverse stresses imposed by host immunity and deploy countermeasures to neutralize immune effectors. The ubiquity of metalloenzymes in all bacteria necessitates acquisition of trace metals such as iron, zinc and manganese, a vulnerability that is exploited by host nutritional immunity, which limits these metals (1). To counter this metal limitation by the host, pathogens deploy diverse high affinity metal acquisition systems including siderophores for iron and zinc (2–4), and transporters and metallophores for zinc and manganese (1, 3, 5–7). In addition to metal limitation, the host also deploys metals as antimicrobial effectors to kill bacteria (8). Copper and zinc are deposited into the phagosome of infected macrophages as antimicrobials and pathogens employ metal resistance systems such as metal efflux transporters (9–11) and metal binding proteins controlled by metal dependent repressors (9–11). Although both metal limitation and metal intoxication limit pathogen growth, in most cases the essential bacterial metalloenzymes rendered dysfunctional by nutritional immunity are incompletely defined.

*M. tuberculosis* is a successful global pathogen that can survive in both macrophages and neutrophils. *M. tuberculosis* experiences both high copper (10–15) and high zinc (12) in the macrophage phagosome and resists metal toxicity through several mechanisms including efflux and metal chelation. Recent data also indicates that *M. tuberculosis* experiences zinc starvation during infection, possibly imposed by calprotectin in caseum (16). However, it is unknown whether copper acquisition, and resistance to copper deprivation, are part of the virulence program of *M. tuberculosis*. *M. tuberculosis* synthesizes diisonitrile lipopeptide natural products directed by the 5 gene *nrp* operon, present in *M. tuberculosis*, *M. bovis*, and *M. marinum* (17–21). The *nrp* operon is induced by copper deprivation and growth of *M. tuberculosis* lacking diisonitrile lipopeptide biosynthesis is inhibited in copper limiting conditions and rescued by a synthetic diisonitrile (22), establishing diisonitrile lipopeptides as mycobacterial chalkophores. However, the role of diisonitrile chalkophores in *M. tuberculosis* pathogenesis is not understood, including whether copper deprivation is imposed by the host during infection, and the specific bacterial pathways that require copper supplied by the chalkophores. In this study, we demonstrate that diisonitrile chalkophores supply copper to the heme-copper *bcc*:*aa_3_* respiratory supercomplex to maintain respiration and ATP production. The heme-copper oxidase of chalkophore deficient *M. tuberculosis* is compromised by the host during infection, but *M. tuberculosis* compensates with the heme-dependent cytochrome BD (CytBD). *M. tuberculosis* lacking diisonitrile chalkophore biosynthesis and CytBD is severely attenuated, demonstrating that this multilayered system for protecting respiration is a critical virulence function of *M. tuberculosis*.

## Results

### Chalkophore deficient *M. tuberculosis* upregulates respiratory chain components in response to copper deprivation

To understand the function of diisonitrile chalkophores in *M. tuberculosis*, we examined the transcriptional profile of wild type (WT) and Δ*nrp M. tuberculosis* treated with 10 μM tetrathiomolybdate (TTM), a copper chelator previously shown to inhibit the growth of *M tuberculosis* lacking chalkophore biosynthesis (22). Copper chelation had relatively few effects on gene expression in wild type *M. tuberculosis* (Figure 1A, Table S3). In contrast, in Δ*nrp M. tuberculosis* we observed upregulation of the genes of the chalkophore cluster (other than *nrp* itself) at baseline, consistent with prior data demonstrating autoregulation (22). In addition, a cluster of genes was induced by copper deprivation in Δ*nrp* cells, but not wild type cells, encoding components of the respiratory chain, including *cydABDC* (encoding components of the heme dependent oxidase CytBD), *qcrABC* (encoding subunits of the *bcc:aa_3_* heme-copper oxidase), and subunits of ATP synthase (Fig 1A). Prior work demonstrated that genetic or pharmacologic disruption of the *bcc:aa_3_* respiratory oxidase, including by Q203, an inhibitor of the QcrB subunit of the *bcc:aa_3_* oxidase, transcriptionally upregulates genes encoding components of the respiratory chain, including *cydAB* and ATP synthase (23, 24). Treatment of wild type and Δ*nrp* with Q203 reproduced the published pattern in wild type cells and indicated that diisonitrile chalkophore deficient cells still respond to Q203 (Figure 1). To confirm the RNA sequencing results, we quantitated the transcript encoding the *cydA* encoded subunit of the CytBD oxidase with different concentrations of TTM and observed no induction at any TTM concentration in wild type cells, but progressive induction of CydA with escalating TTM concentrations in the Δ*nrp* strain (Figure 1B). Longer durations of copper deprivation induced *cydA* in both wild type and Δ*nrp*, with higher induction in Δ*nrp* (Figure 1C). Escalating concentrations of TTM did not affect wild type *M. tuberculosis* growth but caused a graded inhibition of growth of the Δ*nrp* strain in liquid media (Fig 1D). These data indicate that copper deprivation in *M. tuberculosis* lacking diisonitrile chalkophore biosynthesis stimulates gene expression that mimics inhibition of the *bcc:aa_3_* respiratory oxidase.

**Figure 1.**
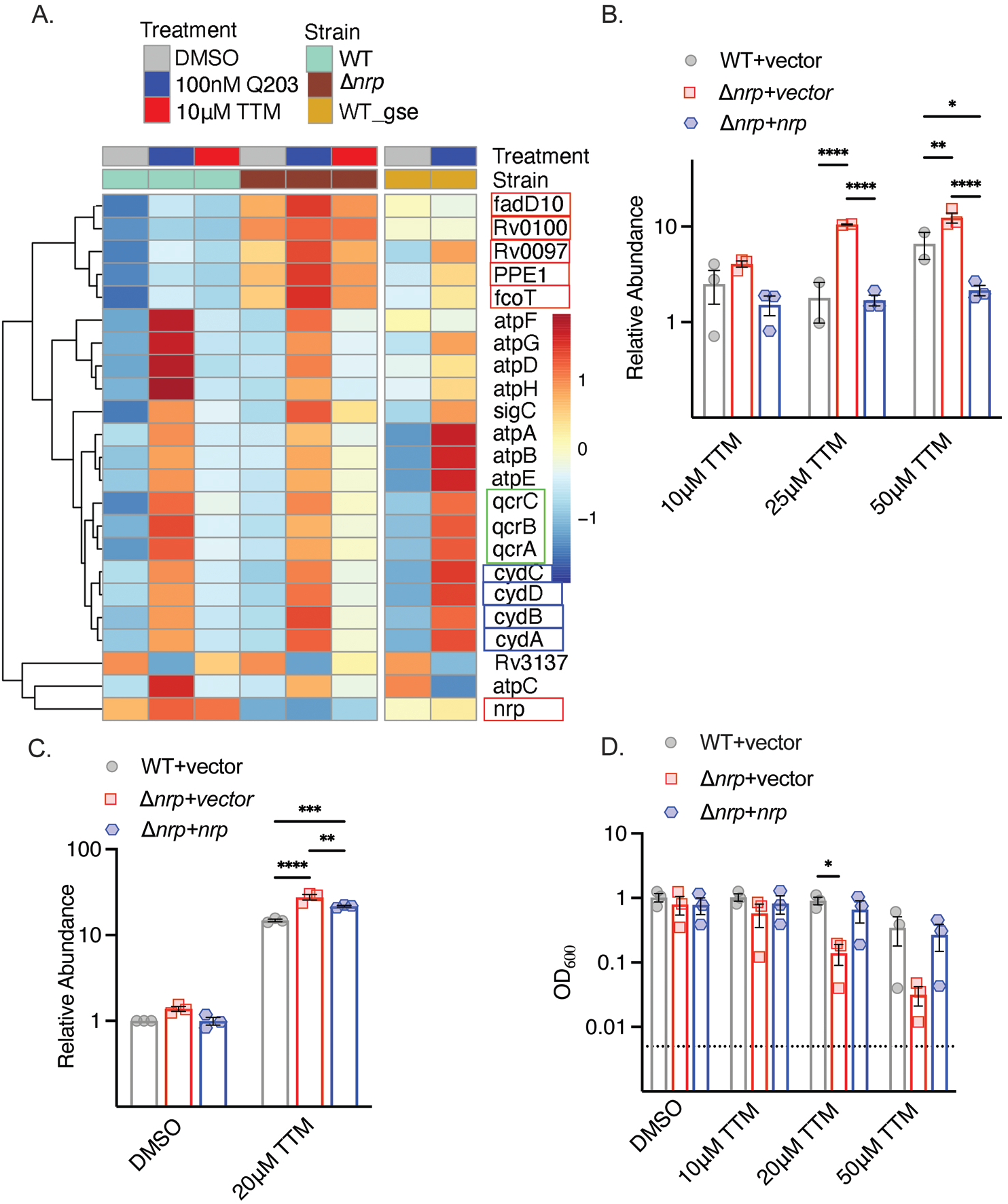
Copper deprivation in chalkophore-deficient *M. tuberculosis* mimics *bcc:aa_3_* oxidase inhibition. **A.** Heat map of transcripts encoding selected respiratory chain components determined by RNA sequencing of *M. tuberculosis* WT or Δ*nrp* treated with TTM or Q203. WT_GSE is the published dataset GSE159080 of *M. tuberculosis* H37Rv treated with Q203. Genes in the chalkophore cluster are boxed in red, genes encoding the CytBD oxidase in in blue, and genes encoding components of the *bcc*:*aa3* supercomplex are in green. **B.** RT-qPCR of the transcript encoding CydA in *M. tuberculosis* WT, Δ*nrp*, and complemented strain treated with varying TTM concentrations for 4 hours. Error bars represent standard error of the mean (SEM). Statistical significance determined via two-way ANOVA with Tukey correction for multiple comparisons. *=p<0.05, **=p<0.01, ****= p<0.0001. **C.** RT-qPCR of the transcript encoding CydA in *M. tuberculosis* WT, Δ*nrp*, and complemented strain treated with 20 μM TTM for 24 hours. Error bars are SEM. Statistical significance determined via two-way ANOVA with Tukey correction for multiple comparisons. **=p<0.01, ***=p<0.001, ****= p<0.0001. **D.** Dose dependent effect of TTM on growth of the indicated *M. tuberculosis* strains at 7 days post inoculation. The dotted line indicates the starting inoculum. Error bars are SEM. Statistical significance determined via two-way ANOVA with Tukey correction for multiple comparisons. *=p<0.05.

### Chalkophores protect the *bcc:aa_3_* oxidase from copper deprivation

Inhibition of *bcc:aa_3_* oxidase by Q203 (23, 25) or other QcrB inhibitors (26–29) is bacteriostatic and incompletely inhibits respiration, but becomes bactericidal and abolishes oxidative phosphorylation in *M. tuberculosis* lacking the alternative cytochrome BD (Δ*cydAB*, (23, 29)) or treated with a CytBD inhibitor (30). Although the *bcc:aa_3_* oxidase and CytBD are functionally redundant, they differ in cofactor usage for electron transfer: *bcc:aa_3_* is a heme-copper oxidase with three copper ions in the electron transport path (31–34), whereas CytBD uses heme prosthetic groups but not copper (35)(Figure 2A).

**Figure 2.**
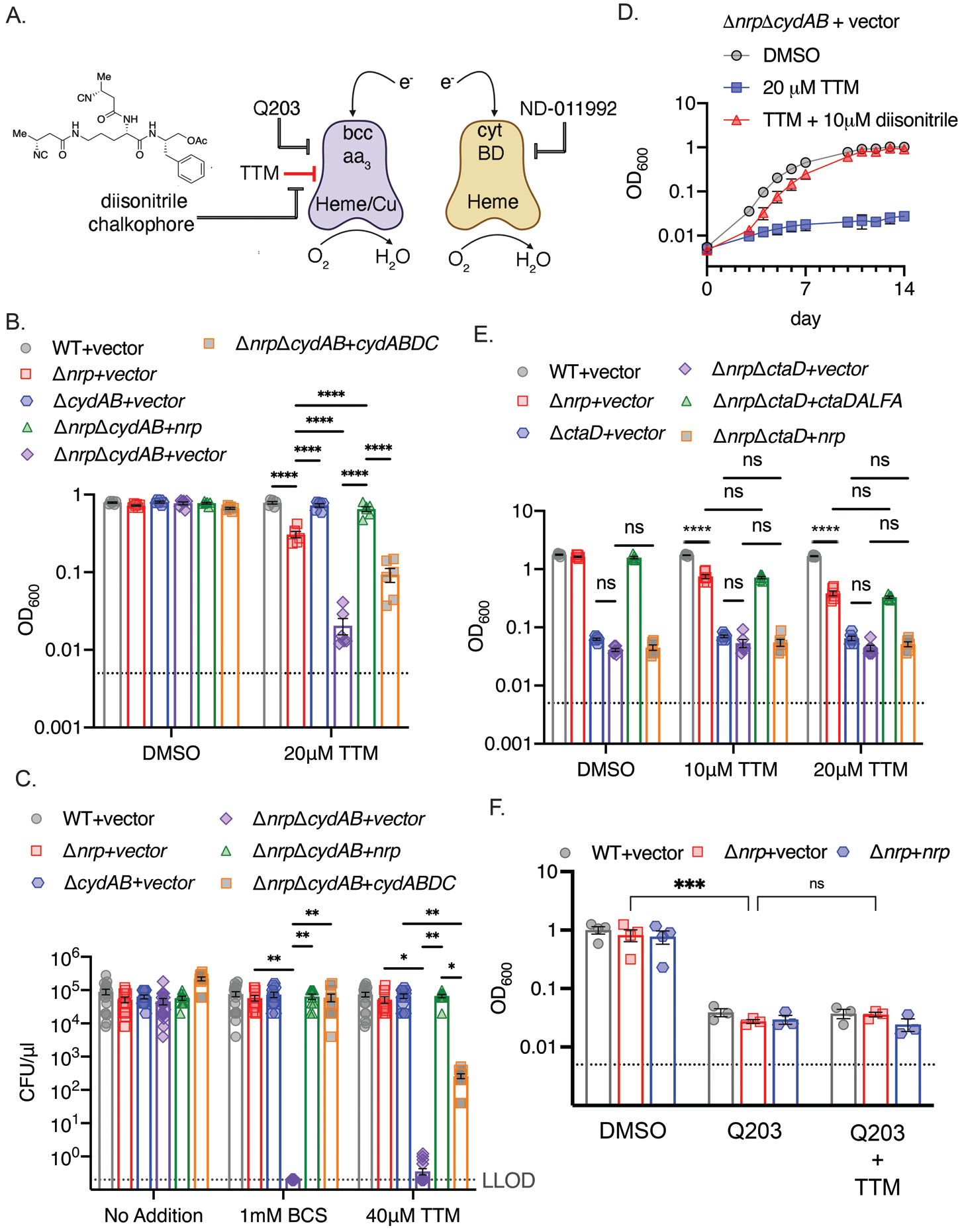
Chalkophores maintain *M. tuberculosis* viability through the heme-copper *bcc:aa_3_*oxidase during copper starvation. **A. Schematic of the terminal respiratory oxidases of *M. tuberculosis*.** The *bcc:aa_3_* oxidase is a heme-copper oxidase and CytBD is a copper-independent heme oxidase. Both transfer electrons to oxygen. Q203 is an inhibitor of *bcc:aa_3_* by targeting the QcrB subunit, whereas ND-011992 targets CytBD. The two oxidases are individually dispensable due to compensation by the other oxidase, but *M. tuberculosis* lacking both is nonviable. The model to be tested is that copper chelation deprives the *bcc:aa_3_* oxidase of copper and that diisonitrile chalkophores counter this copper deprivation stress. **B.** Liquid growth assays of the indicated strains with or without 20 µM TTM treatment. OD_600_ at day 10 post inoculation displayed. Dotted line indicates starting inoculum. Error bars are SEM. Statistical significance determined via two-way ANOVA with Tukey correction for multiple comparisons. ****= p<0.0001. **C.** Bacterial survival of the indicated strains on agar media containing DMSO, 1 mM BCS or 40 μM TTM. Dotted line indicates lower limit of detection (LLOD). Error bars are SEM. Statistical significance determined via two-way ANOVA with Tukey correction for multiple comparisons. **= p<0.01, *=p<0.05 **D. The copper deprivation sensitivity of *M. tuberculosis* Δ*nrp*Δ*cydAB* strain can be rescued with a synthetic diisonitrile chalkophore.** Liquid growth assays of Δ*nrp*Δ*cydAB* with DMSO, 20 μM TTM, or 20 μM TTM with 10 μM of the diisonitrile chalkophore pictured in panel A. Error bars are SEM. **E. The *bcc:aa_3_* oxidase is the only target of copper starvation countered by diisonitrile chalkophores.** Liquid growth assays of the indicated strains treated with 10 or 20 µM TTM, or DMSO vehicle control. OD_600_ at day 10 post inoculation displayed. Dotted line indicates starting inoculum. Error bars are SEM. Statistical significance determined via two-way ANOVA with Tukey correction for multiple comparisons. ns= not significant, ****= p<0.0001. **F. The effect of copper deprivation is masked by inhibition of QcrB subunit of *bcc*:*aa_3_*.** Liquid growth assays of the indicated strains treated with Q203 (100nM) alone or co-treated with 100nM Q203 and 10μM TTM. OD_600_ at day 7 post inoculation displayed. Dotted line indicates starting inoculum. Error bars are SEM. Statistical significance determined via two-way ANOVA with Tukey correction for multiple comparisons. ***=p<0.001, ns= not significant.

To test whether diisonitrile chalkophores maintain the function of the copper dependent *bcc:aa_3_* oxidase under copper limitation (Figure 2A), we generated *M. tuberculosis* Δ*nrp*Δ*cydAB*, along with control strains lacking *cydAB* alone and genetically complemented strains. We tested the effect of TTM on growth in liquid media and observed mild inhibition of growth of *M. tuberculosis* Δ*nrp* and no effect on Δ*cydAB* (Figure 2B). However, growth of Δ*nrp*Δ*cydAB* was severely inhibited by TTM, an effect that was reversed by genetic complementation with either *nrp* or *cydABDC* (Figure 2B). Assays on copper chelated agar media (BCS or TTM) revealed a dramatic sensitization of chalkophore deficient *M. tuberculosis* by loss of the secondary oxidase with 6 logs of killing, a phenotype that was also complemented by *nrp* or *cydABDC* (Figure 2C, S1), although complementation was incomplete with TTM compared to BCS for reasons we have not determined. To determine whether the phenotypes of Δ*nrp* are due to diisonitrile chalkophore deficiency rather than some function of the genetic element independent of the diisonitrile chalkophore itself, we first repeated the same *cydAB* synergy test with *M. tuberculosis* Δ*fadD10*, a second biosynthetic gene in the *nrp* operon that is required for chalkophore mediated resistance to copper chelation (22). Δ*fadD10*Δ*cydAB M. tuberculosis* was also dramatically sensitized to bathocuproinedisulfonic acid, disodium salt (BCS, another copper chelator) or TTM to a similar degree as Δ*nrp*Δ*cydAB* (Figure S1B, C). To demonstrate that the copper deprivation sensitivity of the Δ*nrp*Δ*cydAB* strain is due to absence of the diisonitrile lipopeptide, we synthesized an analogue of a reported *M. tuberculosis* diisonitrile chalkophore (21) having 4-carbon side chains (see Methods and SI Appendix). This C4 l-ornithyl-l-phenylalaninol acetate diisonitrile (C4-Orn-Phin-OAc) was synthesized via a route analogous to our previously reported syntheses of a *Streptomyces* diisonitrile (36). This synthetic diisonitrile efficiently rescued Δ*nrp*Δ*cydAB* from the growth inhibition imposed by TTM in liquid media (Figure 2D), consistent with direct mediation of this effect by the diisonitrile lipopeptide product of the *nrp* locus.

The data above indicates that diisonitrile biosynthesis is necessary to resist the effects of copper starvation and that one of the effects of this copper starvation is dysfunction of the *bcc:aa_3_* oxidase. To determine whether maintenance of respiratory oxidase function under copper starvation is the only function of diisonitrile chalkophores, we executed genetic and biochemical epistasis testing. Deleting *ctaD,* which encodes one subunit of the *bcc:aa_3_* supercomplex, alone or in diisonitrile chalkophore deficient cells (Δ*nrp*Δ*ctaD*), impaired growth of *M. tuberculosis* in copper replete media (Figure 2E), but copper deprivation had no additional effect, indicating that the heme-copper oxidase complex is the only target impacted by copper starvation in these culture conditions (Figure 2E). Similarly, the growth inhibitory effect of copper starvation on Δ*nrp* cells was similar in degree to treatment of WT or Δ*nrp* cells with Q203, but combined treatment was not synergistic (Figure 2F), again indicating that the heme-copper oxidase is inactivated by copper deprivation when the diisonitrile chalkophore is missing and that no additional targets are relevant to copper deprivation-induced growth arrest in these conditions.

### Chalkophores maintain oxidative phosphorylation and ATP production

To determine whether diisonitrile chalkophores directly maintain respiration during copper deprivation, as suggested by the growth and survival data above, we measured oxygen consumption by *M. tuberculosis* using a qualitative methylene blue assay (23, 30). In this assay, decolorization of methylene blue in a sealed tube indicates oxygen depletion, as observed in WT or Δ*nrp*Δ*cydAB M. tuberculosis* treated with DMSO (Figure 3A), indicating intact respiration in chalkophore deficient cells in basal conditions. Treatment with Q203 did not impair oxygen consumption in WT cells but did in Δ*nrp*Δ*cydAB*, consistent with its inhibition of the *bcc*:*aa*_3_ oxidase (Figure 3A). Treatment of wild-type cells with Q203 in combination with ND-011992, a small-molecule inhibitor of CytBD, also blocked oxygen consumption, consistent with prior data (Figure S4A) (23, 30). Copper chelation with 50 μM TTM had no effect on wild type cells but abolished respiration in Δ*nrp*Δ*cydAB*, indicating that copper deprivation inhibits respiration through the *bcc:aa_3_* oxidase (Figure 3A). Similarly, Δ*nrp* cells grown on agar media with TTM and ND-011992 lost viability (Figure S4C). Taken together, these data are consistent with a model in which *M. tuberculosis* can respire via either the *bcc*:*aa_3_* oxidase or CytBD, with the former requiring diisonitrile chalkophore-mediated copper acquisition under copper starvation. The full effect of loss of chalkophore biosynthesis is masked by the redundancy of the oxidases, but in cells that rely only on *bcc*:*aa_3_* oxidase, respiration is abolished by copper deprivation.

**Figure 3.**
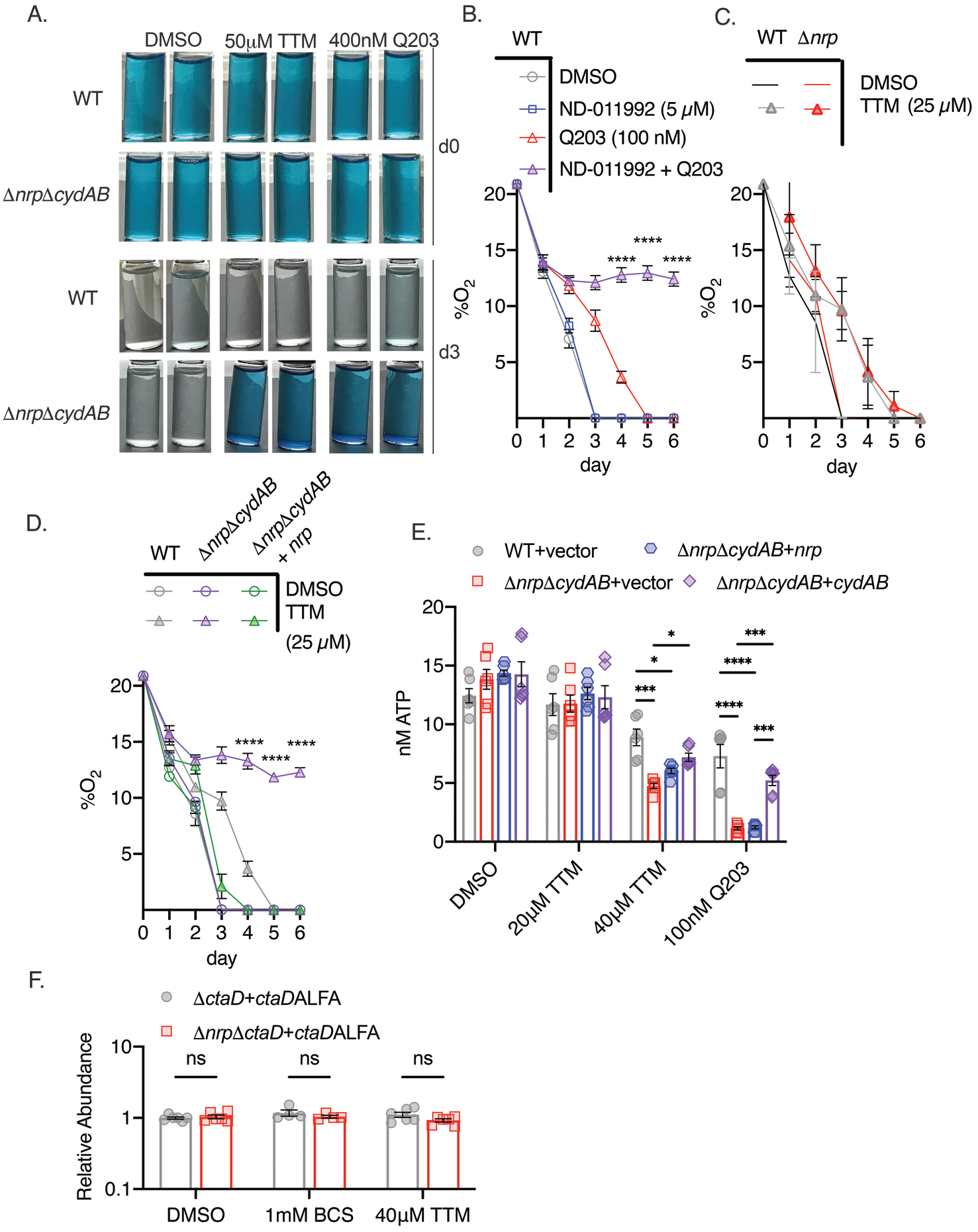
Chalkophore biosynthesis maintains oxidative phosphorylation through the heme-copper *bcc:aa_3_*oxidase. **A.** Methylene blue decolorization assay of oxygen consumption under copper deprivation (TTM) or treatment with Q203 in WT or Δ*nrp*Δ*cydAB M. tuberculosis* at day 0 (d0) or day 3 (d3) of incubation. Clear vials indicate oxygen consumption by respiration. **B.** Quantitative measurement of oxygen consumption using oxygen sensitive optical sensors. WT *M. tuberculosis* treated with DMSO, ND-011992, Q203, or both ND-011992 and Q203. Oxygen measurements were taken daily. Each point represents three measurements of two biologic replicates. Error bars are SEM. Statistical significance between Q203 and ND-011992 + Q203 determined via two-way ANOVA with Tukey correction for multiple comparisons. ****= p<0.0001. **C.** Same assay as in panel B with WT and Δ*nrp M. tuberculosis* treated with DMSO or 25 μM TTM. Error bars are SEM. **D.** Same assay as in panel B with WT, Δ*nrp*Δ*cydAB,* or Δ*nrp*Δ*cydAB* + *nrp* treated with 25 μM TTM. Error bars are SEM. Statistical significance between WT and Δ*nrp*Δ*cydAB* treated with 25 μM TTM determined via two-way ANOVA with Tukey correction for multiple comparisons. ****= p<0.0001. **E.** Cellular ATP levels determined by BacTiter-Glo in the indicated strains treated with DMSO, 20 or 40 μM TTM, or 100 nM Q203. [ATP] determined by standard curve determined in growth media containing the same quantities of DMSO, TTM or Q203. Error bars are SEM. Statistical significance determined via two-way ANOVA with Tukey correction for multiple comparisons. *= p<0.05, ***=p<0.001, ****=p<0.0001. **F.** Relative abundance of a CtaD-ALFA protein in *M. tuberculosis* of the indicated genotype treated with BCS or TTM. See Figure S4 for primary immunoblot data. Error bars are SEM. Statistical significance determined via two-way ANOVA with Tukey correction for multiple comparisons. ns= not significant.

To measure oxygen consumption more quantitatively, we adopted a spot sensor assay (37), that can noninvasively measure oxygen in sealed vessels, allowing serial measurements. WT *M. tuberculosis* consumed oxygen down to the lower limit of detection, and treatment with the CytBD inhibitor ND-011992 (30) had no effect (Figure 3B). Treatment with the QcrB inhibitor Q203 delayed but did not prevent oxygen consumption, consistent with the methylene blue results and prior data (23, 25, 30). However, treatment with the combination of Q203 and ND-011992 completely inhibited oxygen consumption (Figure 3B). Applying this assay to diisonitrile chalkophore deficient strains, we observed that copper deprivation delayed but did not prevent respiration in Δ*nrp* cells (Figure 3C), consistent with compensation by CytBD, but abolished respiration in Δ*nrp*Δ*cydAB* cells, a phenotype that was rescued by genetic complementation with the *nrp* gene (Figure 3D). To confirm that the diisonitrile chalkophore-mediated protection of respiration is accompanied by ATP depletion, we measured ATP levels in WT and chalkophore deficient cells treated with copper deprivation or Q203. Copper deprivation with TTM or treatment with Q203 had minimal effect on wild-type cells (Figure 3E), but inhibited ATP production in the Δ*nrp*Δ*cydAB* strain (Figure 3E). Taken together, these data demonstrate that diisonitrile chalkophores maintain oxidative phosphorylation by the heme-copper respiratory oxidase under copper starvation. Because the electron transport path of the *bcc:aa_3_* oxidase contains three copper sites, these findings are consistent with a model in which these copper sites become dysfunctional when copper is limiting, leading to impaired supercomplex biosynthesis or function.

In eukaryotic copper trafficking disorders in which copper becomes limiting in mitochondria, cytochrome C oxidase becomes unstable, preventing assembly (38). To determine if similar mechanism is operative in mycobacterial cells, we inserted an ALFA epitope tag (39) at the C terminus of CtaD in both WT and Δ*nrp* cells and, after confirming functionality (Figure 2E), examined CtaD protein levels with copper deprivation. CtaD levels remained unchanged with either TTM or BCS (Figure 3F, S4B), indicating that copper deprivation is not affecting oxidase biogenesis and is more likely acting on preexisting respiratory oxidase complexes.

### Chalkophores defend oxidative phosphorylation from nutritional immunity

Respiration has emerged as an attractive target for antimycobacterials, with the ATP synthase inhibitor bedaquiline now a cornerstone of MDR TB treatment (40, 41) and Q203 in clinical trials (42). Although it is also clear that ATP generation via oxidative phosphorylation is required for *M. tuberculosis* growth in mice (23, 30, 43, 44), it is less clear whether the reactive centers of the electron transport chain are compromised by the host and whether *M. tuberculosis* must defend or restore the integrity of the electron transport chain during infection. To examine this question, we infected C57BL/6 mice with *M. tuberculosis* strains lacking diisonitrile chalkophore biosynthesis, alone and in combination with deletion of the genes encoding CytBD (Δ*nrp* and Δ*nrp*Δ*cydAB*). The mouse attenuation phenotype of *M. tuberculosis* lacking *nrp* has been reported by several groups (12, 18, 21) and indicates a mild early attenuation phenotype in the lungs. Loss of *cydAB* in the Δ*nrp* background dramatically exacerbated the mild attenuation of diisonitrile chalkophore deficient strain in the lung (Figure 4A) and caused severe attenuation in the spleen (Figure 4B). Complementation with the *nrp* gene restored virulence to wild-type levels, demonstrating that chalkophore mediated protection of the respiratory chain is a critical virulence function of *M. tuberculosis*.

**Figure 4.**
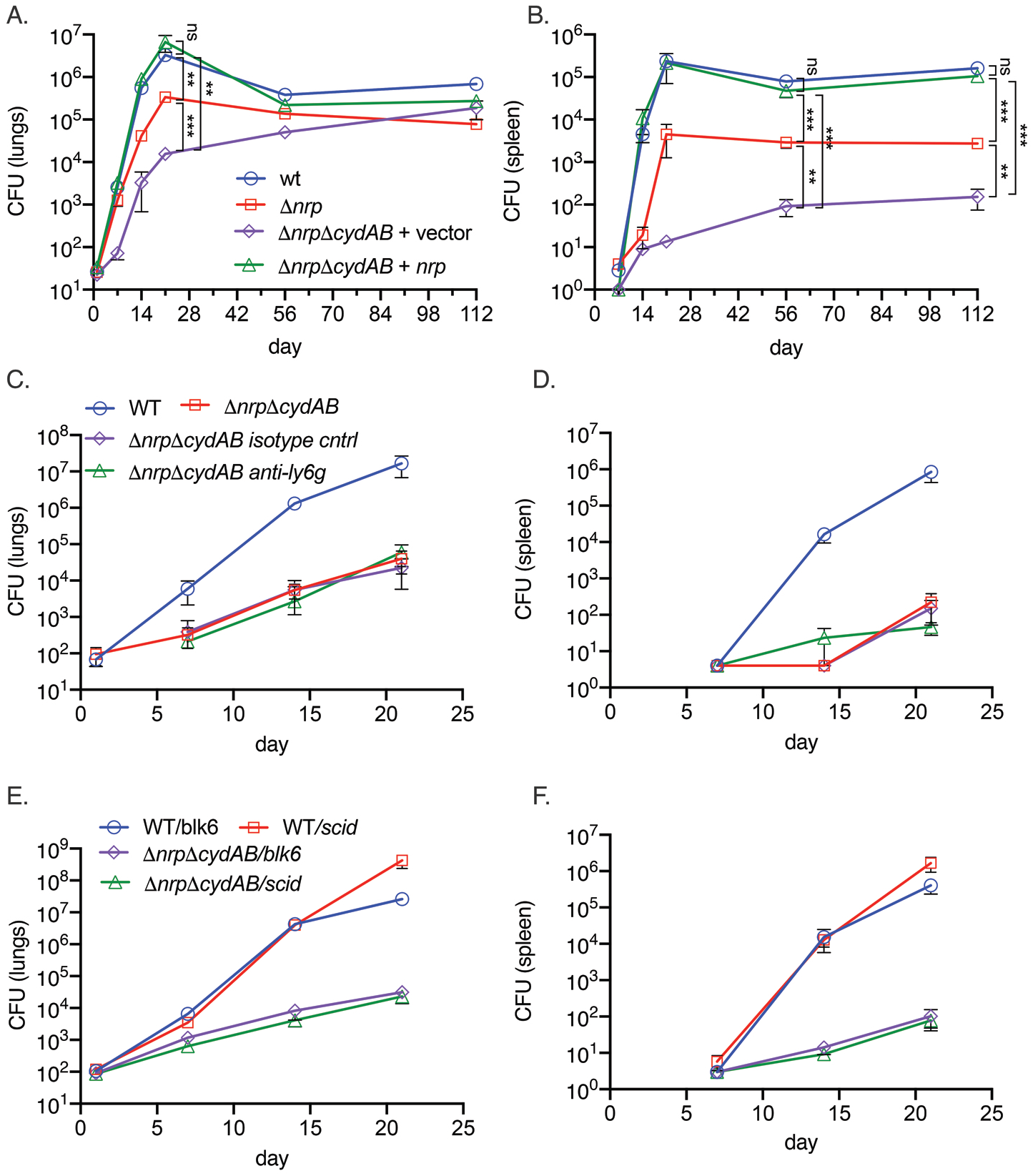
Respiratory chain flexibility is critical for *M. tuberculosis* virulence. A, B. Bacterial titers in the lung (A) or spleen (B) in mice infected with *M. tuberculosis* WT, Δ*nrp*, Δ*nrp*Δ*cydAB,* or Δ*nrp*Δ*cydAB* + *nrp.* Error bars are SEM. Statistical significance determined via two-way ANOVA with Tukey correction for multiple comparisons. Not significant (ns), **=p<0.01, and ***=p<0.001. C, D. **Copper deprivation by the host is independent of neutrophils**. Bacterial titers in the lung (C) or spleen (D) in mice infected with *M. tuberculosis* WT or Δ*nrp*Δ*cydAB,* or Δ*nrp*Δ*cydAB* + *nrp* treated with isotype control antibodies or anti-Ly6G antibodies to deplete neutrophils. Flow cytometric quantitation of neutrophil depletion is provided in Figure S5. Error bars are SEM. E, F. Copper deprivation by the host is independent of adaptive immunity. Bacterial titers in the lung (E) or spleen (F) in C57BL/6J or C57BL/6 SCID mice infected with *M. tuberculosis* WT or Δ*nrp*Δ*cydAB.* Error bars are SEM.

To investigate the host pressure that targets the respiratory chain, we hypothesized that neutrophils, which express several metal binding proteins such as calprotectin, might be relevant. However, efficient depletion of neutrophils by administration of a Ly6G antibody (Figure S5A) did not reverse the severe attenuation of the Δ*nrp*Δ*cydAB* strain in the lungs or spleen (Figure 4C, D). Similarly, infection of SCID mice, which lack adaptive immunity, also had no effect on diisonitrile chalkophore deficient *M. tuberculosis* titers in lung or spleen (Figure 4E, F) and the rapid mortality of SCID mice infected wild type *M. tuberculosis* was not evident in SCID mice infected with Δ*nrp*Δ*cydAB* (Figure S5B), indicating that non adaptive immunity is fully capable of controlling respiratory chain compromised *M. tuberculosis*.

## Discussion

We have identified diisonitrile chalkophore biosynthesis as an *M. tuberculosis* virulence mechanism that defends against copper starvation during infection. Although it is well established that phagosomal pathogens experience both metal excess and metal deprivation in the host, the general paradigm of copper and zinc nutritional immunity is that high phagosomal levels of these metals are deployed as antimicrobial effectors to limit pathogen growth (10, 11, 45–47). Our data indicate that *M. tuberculosis* must also cope with copper deprivation as a host inflicted stress during infection and that *M. tuberculosis* deploys diisonitrile chalkophores to acquire copper in the host. Diisonitrile chalkophores display an extremely high affinity for copper ions (19, 36, 48), and their ability to overcome copper deprivation by strong copper chelators in vitro and host immunity in vivo indicates that they perform an analogous function to bacterial siderophores, which scavenge iron from host iron limitation. Given the abundant data indicating a high copper environment in the macrophage phagosome, the sites and circumstances of the copper deprived niche of *M. tuberculosis* remain to be determined but could include distinct subcellular compartments (i.e., phagosomal vs cytosolic), phases of infection (ie early infection vs post adaptive immunity), or bacterial localization within different lung compartments such as cavities.

Although deprivation of metals from pathogens, including iron and zinc, is a well-recognized mechanism of innate immunity, in most cases the specific bacterial targets that ultimately mediate the antimicrobial function of metal deprivation are not clearly defined and are assumed to be pleiotropic due to the numerous essential metal dependent enzymes in the bacterial cell. However, our data indicates that, in culture, the copper deprivation countered by the diisonitrile chalkophore system targets a single membrane enzyme complex, the *bcc:aa_3_* supercomplex, a heme-copper respiratory oxidase. *M. tuberculosis* deploys two respiratory oxidases, *bcc:aa_3_* and CytBD, which are redundant for bacterial viability. This dual oxidase arrangement provides respiratory chain flexibility during chemical inhibition of each oxidase, and some evidence suggests that respiratory chain flexibility promotes host adaptation (23, 43, 44). The copper centers of respiratory oxidases that participate in electron flow to oxygen are conserved from bacteria to mitochondria and represent a membrane exposed electroreactive center susceptible to damage. We demonstrate that the chalkophore is critical to maintaining the function of the *bcc*:*aa3* supercomplex, both in vitro and in vivo. However, our data does not indicate whether the chalkophore: Cu complex is an intrinsic part of the supercomplex assembly system or simply replenishes the copper pools of the mycobacterial cell. Although chalkophore mediated protection of the *bcc*:*aa_3_* supercomplex is an important virulence function, we cannot exclude the possibility that additional copper dependent enzymes use chalkophore delivered copper during infection. Our data also reveals that *M. tuberculosis* deploys multilayered strategies to maintain oxidative phosphorylation in the face of host immune pressure. In addition, *M. tuberculosis* deploys a backup oxidase, CytBD, which can support *M. tuberculosis* virulence when the *bcc:aa_3_* oxidase is dysfunctional, such as when QcrB is inhibited by Q203. However, the role of this backup oxidase in pathogenesis was unclear. Although CytBD is induced in mouse lung, peaking at 21 days post infection (49), is required for optimal fitness of *M. tuberculosis* in the lung in competition experiments (44), *M. tuberculosis* lacking CytBD is fully virulent (23). Our results reveal the essential role of this compensatory oxidase in virulence, which is only evident when the *bcc:aa_3_* oxidase in inactivated in vivo in *M. tuberculosis* lacking chalkophore biosynthesis, thereby revealing a complex system for maintaining respiration. These studies further strengthen the rationale for targeting CytBD for antibiotic development (35, 50).

The respiratory chain of *M. tuberculosis* has emerged as a promising drug target. Our data indicates that diisonitrile chalkophore biosynthesis, as a mechanism of protection for the respiratory chain during infection, may provide an alternative approach to target this critical energy generating system. Beyond its importance as a drug target, this study identifies a new mechanism of resistance to copper deprivation that, in concert with CytBD, is part of a central virulence strategy of *M. tuberculosis* to protect oxidative phosphorylation during infection.

## Materials and Methods

### Reagents

Middlebrook 7H10 Agar, 7H9 Broth, dextrose, Tween-80, bovine serum albumin (BSA), and UltraPure DNase/RNase -free distilled water were purchased from Fisher Scientific. Ammonium tetrathiomolybdate (TTM), bathocuproinedisulfonic acid, disodium salt (BCS), dimethyl sulfoxide (DMSO), copper sulfate, zinc sulfate, magnesium sulfate, calcium chloride, and ATP were purchased from Millipore Sigma. Biotechnology (BT) grade Chelex 100 resin, sodium form, was purchased from BioRad. Q203 (Telacebec) was purchased from AbMole. ND-011992 was synthesized as previously described (30). The diisonitrile lipopeptide analogue was synthesized and characterized as described below.

### General growth conditions, strains, and DNA manipulations

*M*. *tuberculosis* Erdman WT and mutant stains were grown and maintained in 7H9 media (broth), or on 7H10 (agar) supplemented with 10% OADC (oleic acid, albumin, dextrose, saline), 0.05% glycerol (7H9-OADC/7H10-OADC). Broth cultures were additionally supplemented with 0.02% Tween-80. Chromosomal deletion mutations were generated by specialized transduction utilizing the temperature sensitive phage phAE87. Mutant strains were confirmed by PCR followed by sequencing. For complete strain list with relevant features see **Table S1**. Plasmids utilized in this study were generated using standard molecular techniques and are listed with their features in **Table S2**.

### Growth assays

For liquid growth assays, the indicated strains were pre grown in non-chelated media until reaching an OD_600_ value of ∼1.0. Cells were then collected by centrifugation (3,700 x *g*, 10 min) and washed twice with chelexed phosphate buffered saline with 0.02% Tween-80 (PBS Tween-80). Growth assays were initiated at a calculated OD_600_ of 0.005 by the addition of 1 mL of washed culture at an OD_600_ of 0.05 to 9 mL of replete 7H9-ADS (albumin, dextrose, saline) generated as described previously (22). TTM, BCS, Q203, ND-011992, and/or synthetic diisonitrile chalkophore were then added at the indicated concentration. Growth at 37°C was assayed via daily OD_600_ measurements.

For agar growth assays, cells were pre-grown and washed as indicated above. Washed cells were then normalized to an OD_600_ of 0.1, serially diluted from 10^0^-10^-6^ in PBS Tween-80, and spotted on 7H10-OADC plates containing the indicated concentrations of TTM, BCS, ND-011992 or DMSO vehicle control in triplicate. Plates incubated at 37°C, 5% CO_2_ were imaged after 14 days of incubation to show relative growth, and CFU counts were enumerated after 21 days of incubation.

### RNA preparation

Triplicate 25 mL cultures of strains indicated above were grown in 7H9-ADS until they reached an OD_600_ of 0.3 to 0.5. Cultures were then treated with the indicated concentration of TTM, 100 nM Q203, or a DMSO vehicle control, for 5 hrs. Following a duplicate wash with Chelex treated PBS Tween-80, cells were collected by centrifugation (3,700 × *g*, 10 min), suspended, and resuspended in 1 mL of TRIzol reagent. Cells in 1 mL of TRIzol were mechanically disrupted with zirconia beads via 2×45 sec pulses in a BioSpec Mini24 BeadBeater with 5 min intervening rest periods on ice. Following lysis, beads were removed by centrifugation (20,000 × *g,* 5 min) and total RNA was isolated using the Direct-zol RNA miniprep kit (Zymo Research) as directed by the manufacturer.

### RT-qPCR

DNAse treatment of RNA purified as noted above was performed using the Turbo DNA-free kit (Invitrogen). A total of 500 ng of resulting total RNA was used to synthesize cDNA via random priming utilizing the iScript™ cDNA Synthesis Kit (Biorad). Real-time qPCR was performed on QuantStudio 6 Pro (Applied Biosystems) using iTaq Universal SYBR Green Supermix (Biorad). For *cydA* expression, normalized cycle threshold (CT) was determined relative to the housekeeping gene *sigA*. Reactions with reverse transcriptase were included for each sample and excluded DNA contamination as a source of amplification signal. Relative expression level was calculated using the formula 2^−(CT*cydA*-CT*sigA*)^. Sequences of gene specific primers are as follows: For *cydA,* 5’-GTCATCGAAGTGCCCTATGT-3’ and 5’- CTGGTATTCCTGCTGCAGAT-3’, and for *sigA,* 5’- CGTCTTCATCCCAGACGAAAT-3’ and 5’-CGACGAAGACCACGAAGAC-3’.

### RNA Sequencing and Data Analysis

After RiboGreen quantification and quality control by Agilent BioAnalyzer, 500 ng of total RNA underwent ribosomal depletion with the NEBNext rRNA Depletion Kit (Bacteria) (NEB catalog # E7850) and library preparation with the TruSeq Stranded Total RNA LT Kit (Illumina catalog # RS-122-1202) according to instructions provided by the manufacturer with 8 cycles of PCR. Samples were barcoded and run on a NovaSeq 6000 in a PE100 run, using the NovaSeq 6000 S4 Reagent Kit (200 Cycles) (Illumina). On average, 27 million paired reads were generated per sample. Post-run demultiplexing and adapter removal were performed and fastq files were inspected using fastqc (51). Trimmed fastq files were then aligned to the reference genome (*M. tuberculosis* H37Rv; NC_000962.3) using bwa mem (52). Bam files were sorted and merged using samtools (53) and gene counts were obtained using featureCounts from the Bioconductor Rsubread package (54). Differentially expressed genes were identified using the DESeq2 R package (55) and subsequent analysis of gene expression was performed in (56, 57).

### Immunoblotting

Duplicate 20 mL cultures of chelated 7H9-ADS replete for all ions were inoculated at an OD_600_ of 0.02. Upon reaching an OD_600_ of 0.6-0.7, cultures were treated with the indicated concentration of TTM, or DMSO vehicle control, for 24 hrs. Cells were collected by centrifugation (3700 x *g*, 10 min) washed once with 1mL of lysis buffer (350 mM sodium chloride, 20 mM Tris pH 8.0, 1 mM 2-mercaptoethanol) prior to suspension in 0.8 mL of lysis buffer plus ∼100μL of zirconia beads. Lysis was performed by 3×45 sec pulses in a BioSpec Mini24 beadbeater with 5 min intervening rest periods on ice. Beads and debris were removed by centrifugation at 20,000 x *g* for 15 min at 4°C, the resulting supernatant was mixed 1:1 with 2x Laemmli sample buffer supplemented with 0.1 M dithiothreitol (DTT). 20 μL of each sample, heated for 10 min at 100°C, was then separated on 4-12% NuPAGE Bis-Tris poly acrylamide gels. Separated proteins were transferred to nitrocellulose and probed with the indicated antibodies. Antibodies used in this study are monoclonal anti-*E. coli* RNA -polymerase β (BioLegend), sdAb anti-ALFA tag-HRP (NanoTag Biotechnologies). Chemiluminescence was visualized using SuperSignal^TM^West Pico Plus chemiluminescent substrate (Thermo Scientific). Blots were imaged on an iBright FL1000 imager (Themo Fisher Scientific). CtaD ALFA blots were quantitated using ImageJ software. Relative ALFA signal was normalized to corresponding Rpoβ loading control levels.

### Methylene blue assay

Cultures were grown in 7H9-ADS to an OD_600_ of 0.5-0.7, washed twice in Chelex treated PBS Tween-80, and adjusted to an OD_600_ of 0.15 with fresh 7H9-ADS and incubated with indicated concentrations of TTM, Q203, ND-011992, or DMSO for 4 hrs. Following incubation, 4-ml screw-cap glass vials were filled with the cultures and methylene blue dye was added at a final concentration of 0.001%. All vials were sealed tightly using a PTFE/rubber seal (Thermo Scientific) and incubated at 37°C in an anaerobic container (Benton Dickinson GasPak EZ container systems) for 3 days.

### Oxygen Consumption Measurements

Oxygen consumption was assessed using PSt6 sensor spots and a Fibox 4 trace instrument (PreSens Precision Sensing GmbH Am BioPark, Germany)(37). Screw cap glass vials (Thermo Scientific) containing the sensor spot PSt6 were filled with cultures grown in 7H9-ADS at a calculated OD_600_ of 0.005 and the indicated concentrations of TTM, Q203, ND-011992, or DMSO. The vials were tightly sealed using a PTFE/rubber seal (Thermo Scientific) and incubated at 37°C in an anaerobic pouch (Benton Dickinson GasPak EZ pouch systems). Percent oxygen was measured daily via the Fibox 4 trace oxygen meter without unsealing the anaerobic environment.

### ATP quantitation

ATP levels were quantified with the BacTiter-Glo^TM^ (Promega). The indicated strains were first pre-grown to an OD_600_ of 0.5-1.0 in 7H9-OADC. Cells were then washed twice with PBS Tween-80 and suspended in 7H9-OADC at an OD_600_ of 0.05. 96-Well plates were inoculated with 100 μL of washed culture in replicate wells. The indicated concentrations of TTM, Q203, or DMSO vehicle control were then added, and plates were incubated at 37°C, 0.05% CO_2_ for 24 hrs. 100 μL of BacTiter-Glo was then added to each well and plates were incubated 15 additional mins at 37°C, 5% CO_2_ prior to reading on a SpectraMax M3 plate reader. [ATP] in test samples was quantified against standard curves of ATP.

### Synthesis of diisonitrile chalkophores

Details of synthesis and characterization are provided in SI Appendix 1.

### Aerosol infection of mice

8–10-week-old C57BL/6J (JAX stock number 00064) and 8-10 week B6.Cg-*Prkdc^scid^/SzJ* (stock number 001913, RRID: IMSR_JAX:001913) were purchased from The Jackson Laboratory. All purchased mice were rested within our animal facility to normalize microbiota for 2 weeks. Care, housing, and experimentation on laboratory mice were performed in accordance with the National Institute of Heath guidelines, and the approval of the Memorial Sloan Kettering Institutional Animal Care and Use Committee (IACUC). Strains for infection were grown to an OD_600_ of 0.5-0.7 in 7H9-OADC, washed twice with PBS Tween-80 followed by brief sonication to disrupt aggregates. Final inoculums were prepared by suspending 8×10^7^ CFU in 10 mL of sterile water. Mice were exposed to 4×10^7^ CFU in a Glas-Col aerosol exposure unit, a dose calibrated to deliver 10-50 CFU per mouse. At the indicated time points post infection, 5 individual mice were humanely euthanized and both lungs and spleens were harvested for CFU determination. Organ homogenates were cultured on 7H10-OADC plates, and CFU was enumerated after 28 days of incubation at 37°C, 5% CO_2_. 20 additional SCID mice (10: WT *M. tuberculosis*, 10: Δ*nrp*Δ*cydAB*) infected as described above, were used to assess survival. Infected mice were monitored daily and humanely euthanized upon frank signs of morbidity.

### Neutrophil depletion

Mice were treated intraperitoneally with anti-Ly6G (BioXcell, clone 1A8, RRID AB_1107721) or isotype control (BioXcell, clone 2A3, RRID AB_1107769) every 48 hrs. Lungs were harvested aseptically and single cell suspensions prepared by homogenization in a Bullet Blender with 2.0 mM zirconium oxide beads for 3 min followed by 100-micron filtration. After lysis of red blood cells with Gibco ACK lysing buffer and blocking of Fc receptors (Invitrogen CD16/CD32 clone 93, RRID AB_467133), cells were stained with live/dead aqua (Thermo) and antibodies to CD45 (FITC, clone 30-F11, BD Biosciences, RRID AB_394609), CD11b (BUV661, clone M1/70, BD Biosciences), CD11c (APC-R700, clone N418, RRID AB_2744277), Ly6C (BV605, clone AL-21, BD Biosciences, AB_2737949), CD80 (PerCP Cy5.5, clone 16-10A1 AB_1727514), CD86 (Pacific Blue, clone GL-1, BioLegend, RRID AB_493466), MHC II (BUV395, clone 2G9, BD Biosciences RRID AB_2741827), GR1 (PE, clone RB6-8C5, BD Biosciences, RRID AB_398532), and Ly6G (APC, clone 1A8, BD Biosciences, RRID AB_394208) and analyzed by flow cytometry on a BD Fortessa cytometer after paraformaldehyde fixation.

## Supporting information

RNA seqeuncing count table

Merged SI: Figures S1-S5 and chemical synthesis appendix

## Acknowledgements

This work was supported by R01AI138446 and P30 CA008748. We acknowledge the use of the Integrated Genomics Operation Core, funded by the NCI Cancer Center Support Grant (CCSG, P30 CA08748), Cycle for Survival, and the Marie-Josée and Henry R. Kravis Center for Molecular Oncology. We thank Christina Stallings for advice about neutrophil depletion and George Sukenick and Rui Wang (MSK) for expert NMR and mass spectral support.

## Author Contributions

J.A.B., Y.O., and M.S.G. conceived and planned the experiments. C.H. synthesized the diisonitrile analogue under the supervision of D.T. J.A.B., Y.O., and M.S.G. carried out the experiments with support and technical expertise from A.B. for neutrophil depletion. J.A.B., Y.O., C.H., D.T., and M.S.G. contributed to the interpretation of the results. M.S.G. took the lead in writing the manuscript. All authors provided critical feedback and helped shape the research, analysis, and manuscript.

