## Supplementary material for "Chalkophore mediated respiratory oxidase flexibility controls *M. tuberculosis* virulence": Merged SI: Figures S1-S5 and chemical synthesis appendix

**Buglino et al**

**Supplementary Materials**

Figures S1-S5

SI appendix 1: chemical synthesis methods

**Figure S1 (related to Figure 2). Loss of *cydAB* severely sensitizes chalkophore deficient *M. tuberculosis* to copper chelation**

**A, B.** Images of triplicate 10-fold serial dilutions of the indicated *M. tuberculosis* strains on agar media containing no chelator, 1 mM BCS, or 40 μM TTM.

**C.** Quantitation of the agar survival assay in panel B. Error bars are SEM. Statistical significance determined by two-way ANOVA with Tukey correction for multiple comparisons. *=$p<0.05$, **=$p<0.01$, ***=$p<0.001$

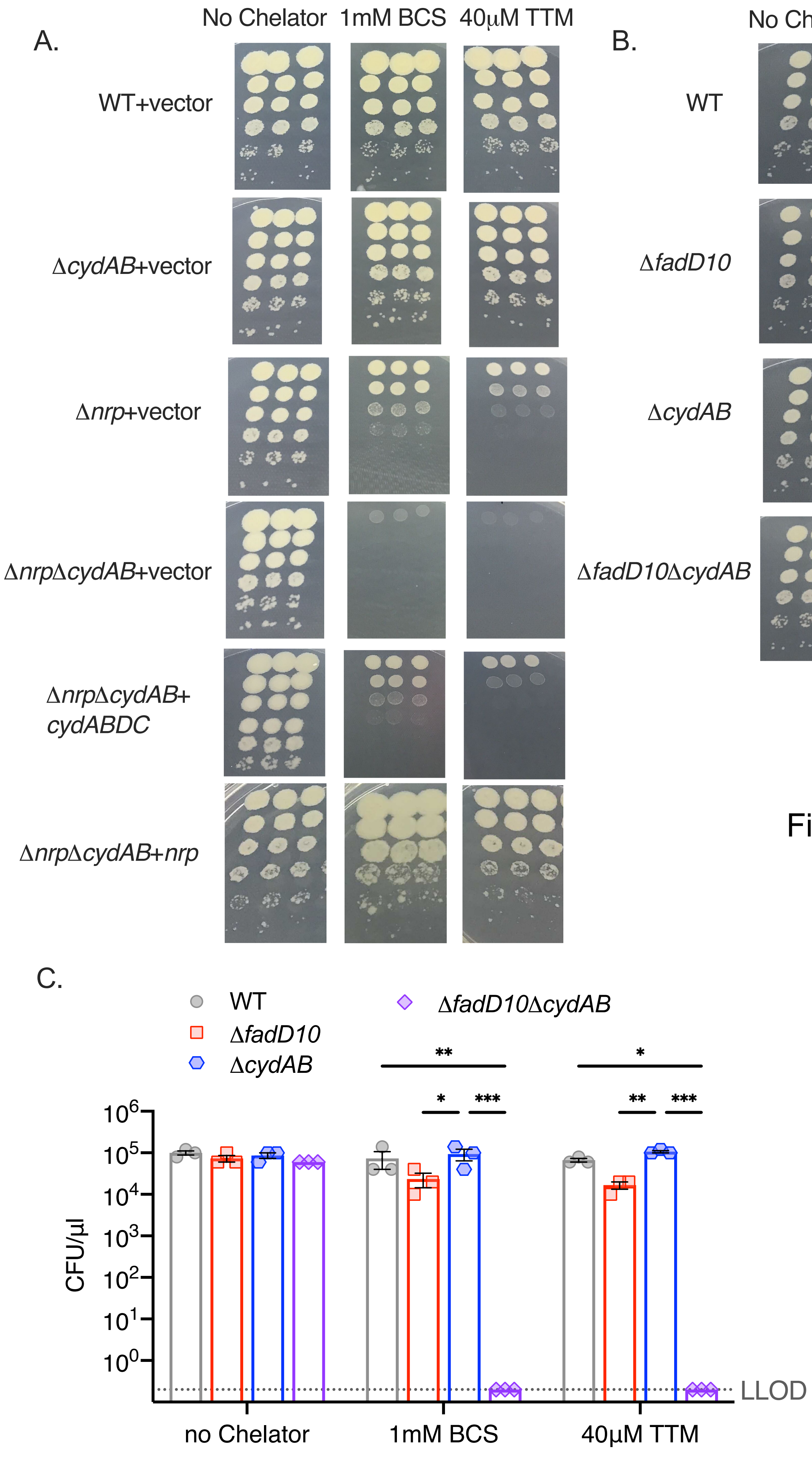
A.
No Chelator
1mM BCS
40μM TTM
WT+vector
ΔcydAB+vector
Δnrp+vector
ΔnrpΔcydAB+vector
ΔnrpΔcydAB+
cydABDC
ΔnrpΔcydAB+nrp
B.
No Chelator
1mM BCS
40μM TTM
WT
ΔfadD10
ΔcydAB
ΔfadD10ΔcydAB
C.
WT
ΔfadD10
ΔcydAB
ΔfadD10ΔcydAB
**
*
*
***
**
***
10^6
10^5
10^4
10^3
10^2
10^1
10^0
CFU/μl
LLOD
no Chelator
1mM BCS
40μM TTM

Figure S1

**Figure S2 (related to Figure 2).**

**A.** Quantitation of optical density in liquid culture from Figure 2D at day 10 time point of the Δ*nrp*Δ*cydAB* strain treated with DMSO, TTM, or TTM+ synthetic diisonitrile. Error bars are SEM. Statistical significance determined by two-way ANOVA with Tukey correction for multiple comparisons ***=$p<0.001$. Dotted line indicates starting inoculum.

**B-F.** Full growth curves in liquid media quantitated in **Figure 2B** for the indicated strains treated with DMSO (grey symbols) or 20 μM TTM (blue symbols). Error bars are SEM and if not visible are within the graphed symbol.

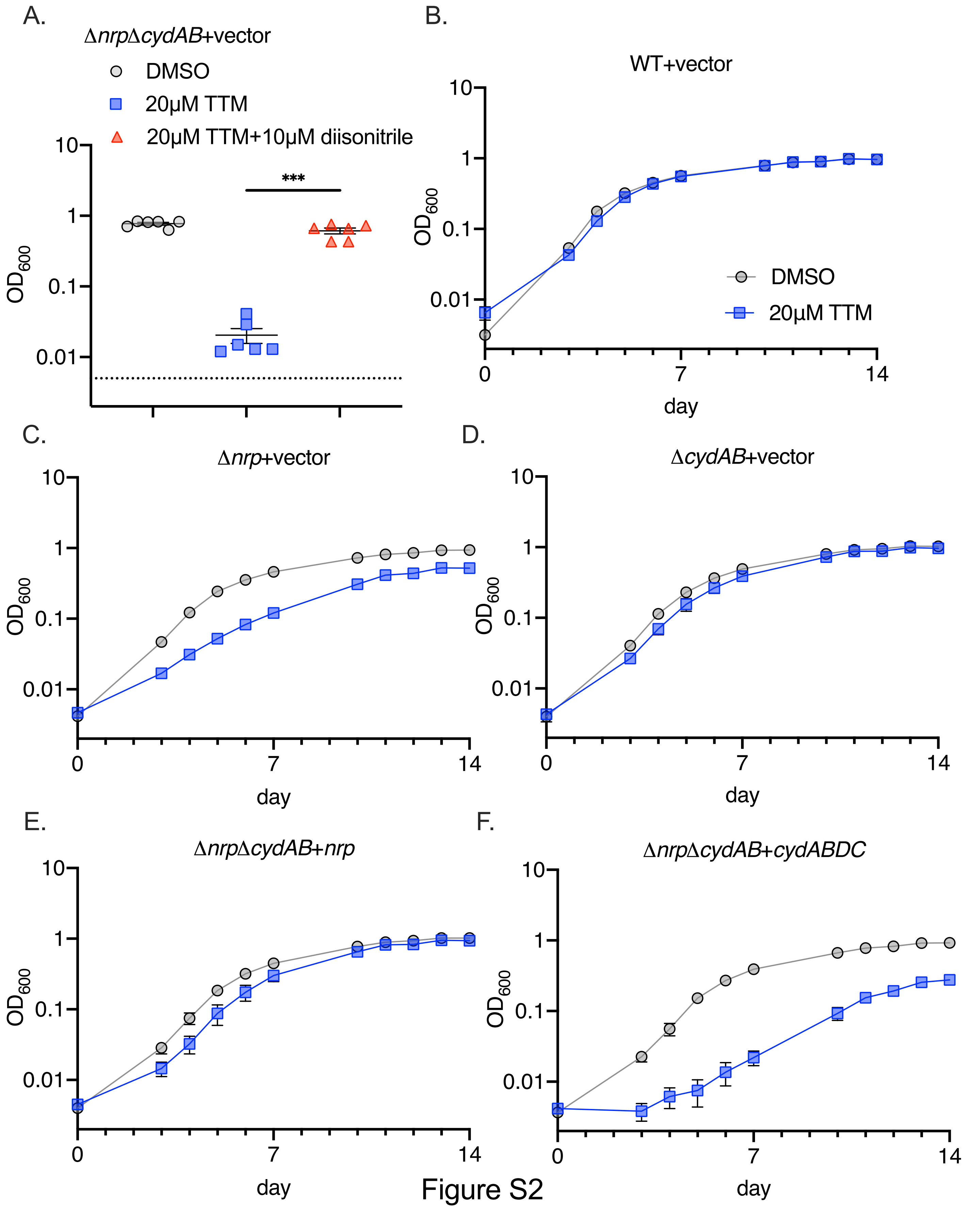

A.
ΔnrpΔcydAB+vector
DMSO
20μM TTM
20μM TTM+10μM diisonitrile
***
OD600
10
1
0.1
0.01
B.
WT+vector
OD600
DMSO
20μM TTM
day
0
7
14
C.
Δnrp+vector
OD600
day
D.
ΔcydAB+vector
OD600
day
E.
ΔnrpΔcydAB+nrp
OD600
day
F.
ΔnrpΔcydAB+cydABDC
OD600
day


Figure S2

**Figure S3 (related to Figure 2E). The *bcc:aa3* oxidase is the only target of chalkophore mediated protection from copper starvation**

**A-B.** Full growth curves in liquid media quantitated in Figure 2E for the indicated strains treated with DMSO (A) or 20 μM TTM (B). Error bars are SEM and are within the symbol if not visible.

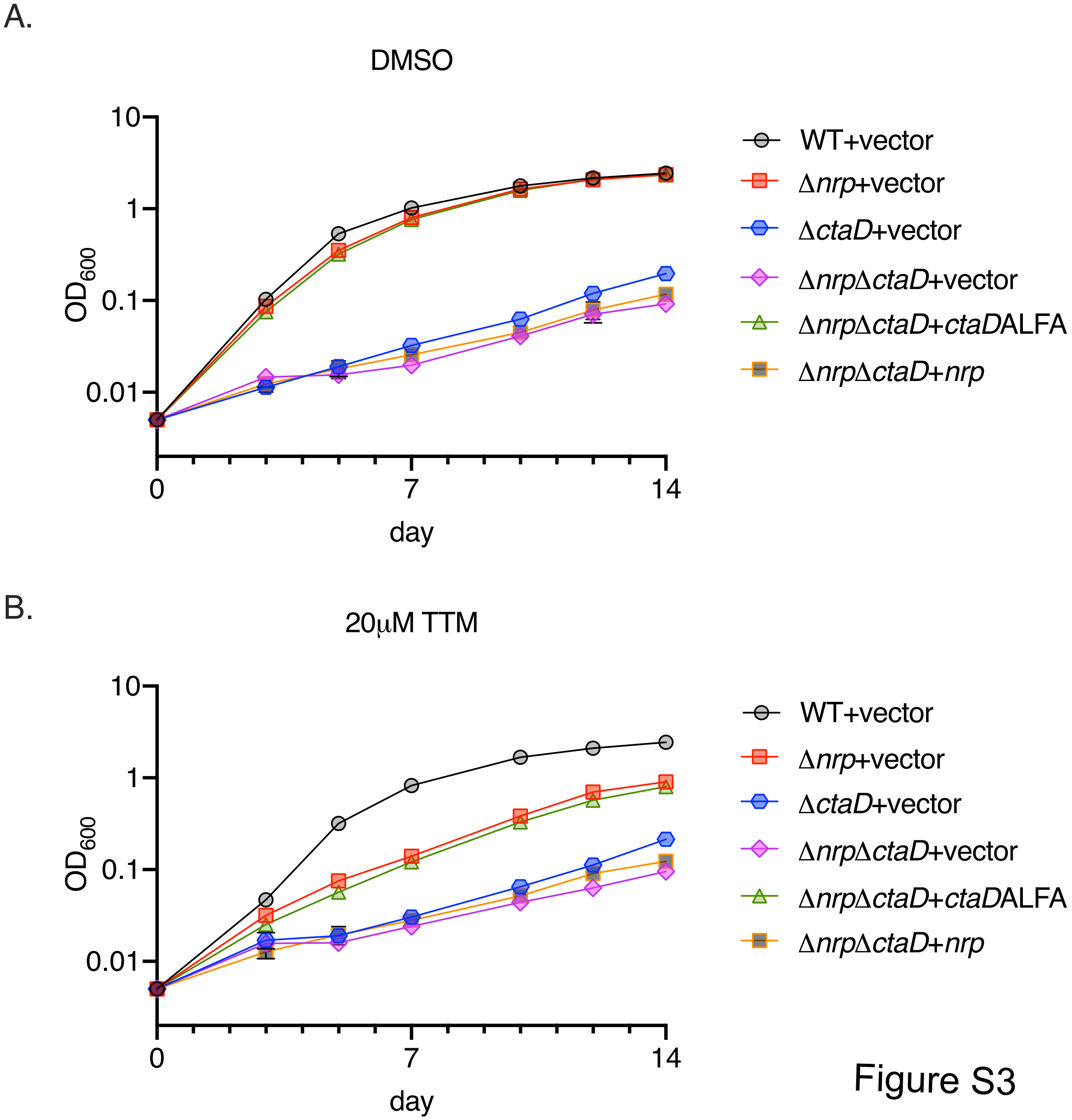
A.
DMSO
10
1
0.1
0.01
OD600
0
7
14
day
WT+vector
Δnrp+vector
ΔctaD+vector
ΔnrpΔctaD+vector
ΔnrpΔctaD+ctaDALFA
ΔnrpΔctaD+nrp
B.
20μM TTM
10
1
0.1
0.01
OD600
0
7
14
day
WT+vector
Δnrp+vector
ΔctaD+vector
ΔnrpΔctaD+vector
ΔnrpΔctaD+ctaDALFA
ΔnrpΔctaD+nrp

Figure S3

**Figure S4 (related to Figure 3). Chalkophores protect the heme-copper respiratory oxidase in copper limiting conditions.**

**A.** Methylene blue decolorization assay in wild type *M. tuberculosis* or *M. tuberculosis* Δ*nrp* at assay start (d0) or after 3 days of incubation in a sealed tube (d3) treated with DMSO, 50 μM TTM, or the combination of Q203 (400 nM) and ND-011992 (50 μM).

**B. Copper chelation does not destabilize the CtaD protein**

*M. tuberculosis* lacking the CtaD subunit of the *bcc:aa3* oxidase and complemented with a fully functional CtaD with a C terminal ALFA epitope tag (see Figure 2E) either in the wild-type background (WT) or *M. tuberculosis* Δ*nrp* (KO), which lacks diisonitrile chalkophore biosynthesis, and treated with either BCS or TTM. Full immunoblots for the ALFA tag or RpoB as a loading control are shown. Images were quantified using ImageJ software.

**C. Copper chelation synergizes with CytBD inhibition in the absence of chalkophore biosynthesis.** Serial dilutions of *M. tuberculosis* WT, Δ*nrp*, Δ*nrp*+*nrp* strains were cultured on agar media containing TTM, ND-011992 (ND), or both. *p=0.0148 by two-way anova.

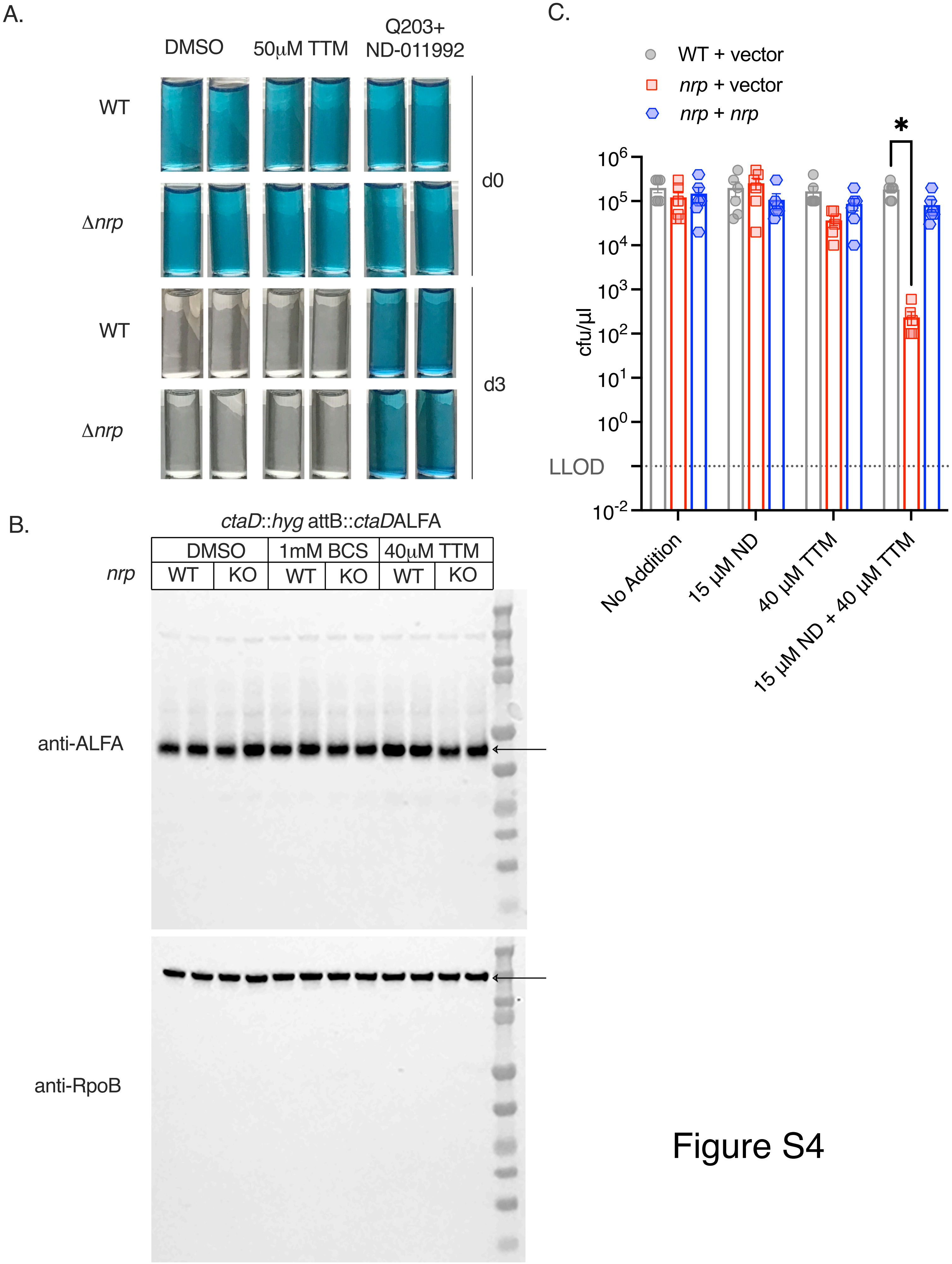
A.
DMSO
50μM TTM
Q203+
ND-011992
WT
Δnrp
WT
Δnrp
d0
d3
B.
ctaD::hyg attB::ctaDALFA
DMSO
1mM BCS
40μM TTM
nrp
WT
KO
WT
KO
WT
KO
anti-ALFA
anti-RpoB
C.
WT + vector
nrp + vector
nrp + nrp
*
cfu/μl
10^6
10^5
10^4
10^3
10^2
10^1
10^0
LLOD
10^-2
No Addition
15 μM ND
40 μM TTM
15 μM ND + 40 μM TTM

Figure S4

**Figure S5 (related to Figure 4) Attenuation of chalkophore-deficient *M. tuberculosis* is independent of neutrophils and adaptive immunity.**

**A.** Flow cytometric quantitation of neutrophils in the lung at day 7 and 14 post infection with the indicated *M. tuberculosis* strains (WT or Δ*nrp*Δ*cydAB*) and treated with isotype control antibody or anti-Ly6G. Neutrophils were defined as $CD11B^{+}GR1^{+}$ and their percentage among CD45+ cells is graphed. Statistical significance determined by unpaired Welch's t test. *=p<0.05, **=p<0.01. Error bars are SEM

**B.** Survival curve of C57BL/6 SCID mice infected with WT or Δ*nrp*Δ*cydAB M. tuberculosis*.

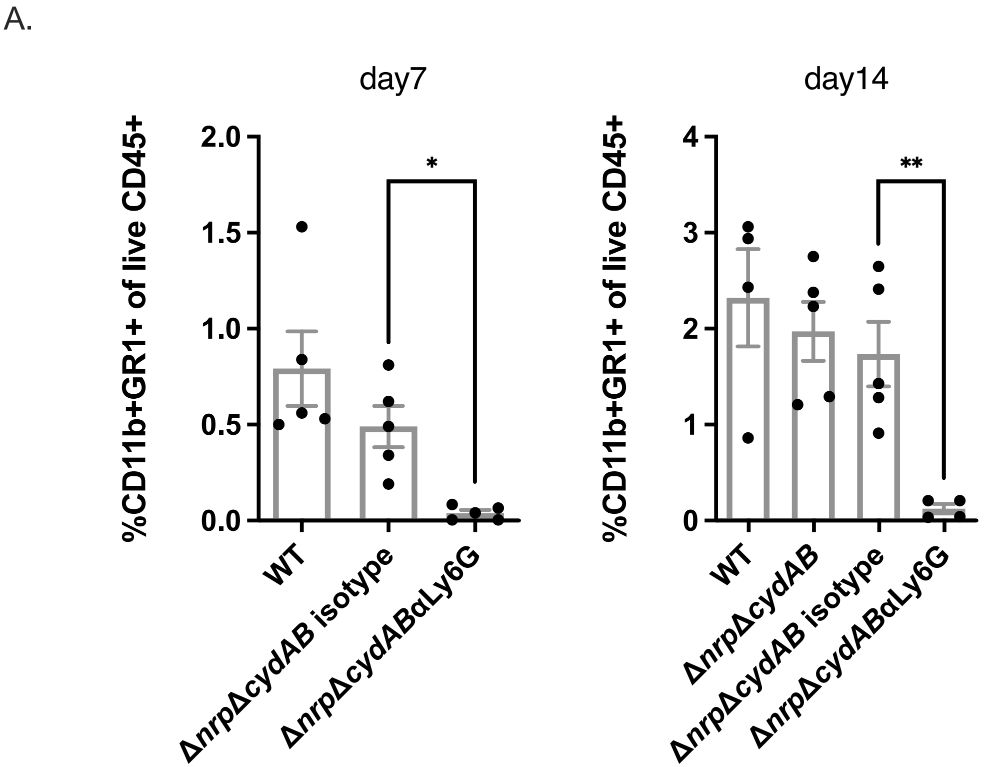
A.
day7
%CD11b+GR1+ of live CD45+
2.0
1.5
1.0
0.5
0.0
*
WT
ΔnrpΔcydAB isotype
ΔnrpΔcydABαLy6G
day14
%CD11b+GR1+ of live CD45+
4
3
2
1
0
**
WT
ΔnrpΔcydAB
ΔnrpΔcydAB isotype
ΔnrpΔcydABαLy6G

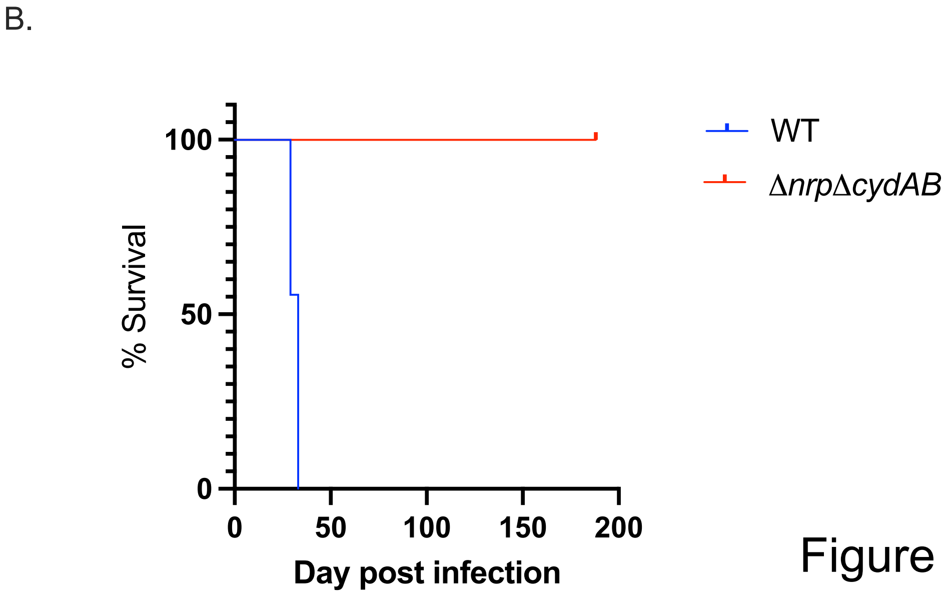
B.
% Survival
100
50
0
0
50
100
150
200
Day post infection
WT
ΔnrpΔcydAB

Figure S5

### Supplemental Appendix I: Chemical Synthesis methods

Buglino et al

#### Supporting Information

A. Synthetic Methods S2
B. Synthesis of Diisonitrile **6** S3
C. $^{1}$H-NMR and $^{13}$C-NMR Spectra S7

#### A. MATERIALS AND METHODS

##### *Reagents*

Reagents were obtained from Aldrich Chemical (www.sigma-aldrich.com) or Acros Organics (www.fishersci.com) and used without further purification. $N^2$,$N^5$-Bis(tert-butoxycarbonyl)-L-ornithine and L-phenylalaninol were obtained from AmBeed (www.ambeed.com) and used without further purification. Optima or HPLC grade solvents were obtained from Fisher Scientific (www.fishersci.com), degassed with Ar, and purified on a solvent drying system as described[1] unless otherwise indicated.

##### *Reactions*

All reactions were performed in flame-dried glassware under positive Ar pressure with magnetic stirring unless otherwise noted. Liquid reagents and solutions were transferred through rubber septa via syringes flushed with Ar prior to use. Cold baths were generated as follows: 0 °C, wet ice/water.

##### *Chromatography*

TLC was performed on 0.25 mm E. Merck silica gel 60 F254 plates and visualized under UV light (254 nm) or by staining with basic potassium permanganate ($KMnO_4$), cerium ammonium molybdenate (CAM), or *p*-anisaldehyde. Silica flash chromatography was performed on E. Merck 230–400 mesh silica gel 60. Parallel chromatography was performed on an ISCO CombiFlash OptiX 10 instrument with RediSep silica gel normal phase columns with ELSD detection and UV detection at 254 nm.

##### *Analytical Instrumentation*

NMR spectra were recorded on a Bruker UltraShield Plus 600 MHz Avance III NMR with DCH CryoProbe at 24 °C in $CDCl_3$ unless otherwise indicated. Chemical shifts are expressed in ppm relative to TMS ($^1$H, 0 ppm) or solvent signals: $CDCl_3$ ($^1$H, 7.26 ppm; $^{13}$C, 77.0 ppm), DMSO-$d_6$ ($^1$H, 2.50 ppm; $^{13}$C, 39.52 ppm); coupling constants are expressed in Hz. NMR spectra were processed using Mnova (www.mestrelab.com/software/mnova-nmr). High resolution mass spectra were obtained on a Waters Acuity Premiere XE TOF LC-MS by electrospray ionization (ESI).

---

[1] Pangborn, A. B.; Giardello, M. A.; Grubbs, R. H.; Rosen, R. K.; Timmers, F. J. *Organometallics* **1996,** *15*, 1518–1520.

#### A. Synthesis of Diisonitrile 6

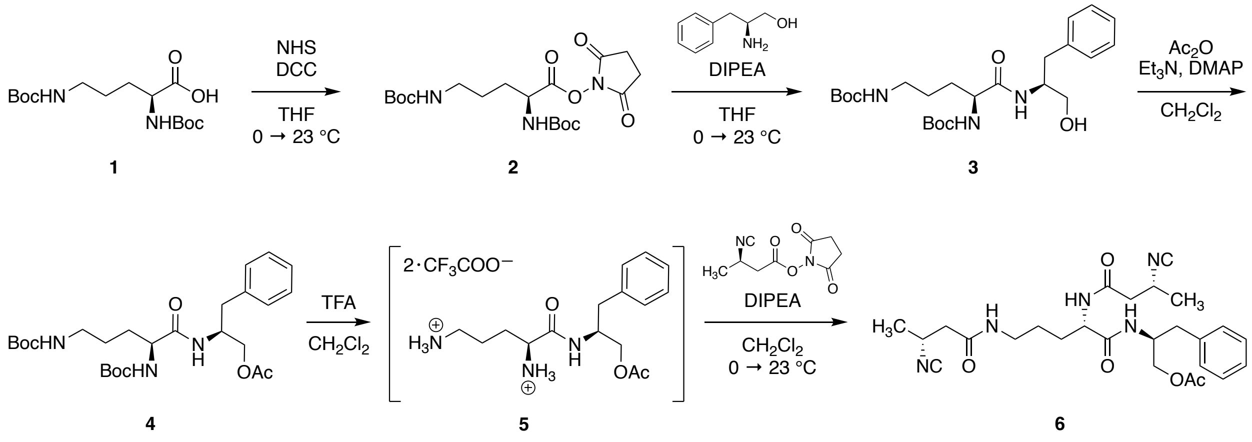


**Synthesis of diisonitrile 6.** Boc = *t*-butyloxycarbonyl; DIPEA = diisopropylethylamine; DMAP = 4-dimethylaminopyridine; NHS = *N*-hydroxysuccinimide; TFA = trifluoroacetic acid; THF = tetrahydrofuran.

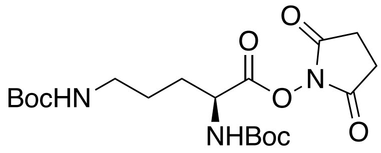


***N*$^2$,*N*$^5$-Bis(*tert*-butoxycarbonyl)-L-ornithine *N*-hydroxysuccinimide ester (2, Boc-Orn[Boc]-NHS).** In a 50-mL roundbottom flask, *N*$^2$,*N*$^5$-bis(*tert*-butoxycarbonyl)-L-ornithine (**1**) (2.0099 g, 6.04 mmol, 1.0 equiv) and dicyclohexylcarbodiimide (1.2932 g, 6.26 mmol, 1.03 equiv) were suspended in 20 mL anhyd THF and stirred at 0 °C for 10 min. *N*-Hydroxysuccinimide (699 mg, 6.07 mmol, 1.005 equiv) was added to the stirring solution in one portion. The mixture was stirred at 0 °C for 30 min, warmed to 22 °C, and stirred for 23 h. The mixture was filtered through a pad of celite and the solid white residue was washed with $Et_2O$ (40 mL). The combined filtrates were concentrated by rotary evaporation to afford the crude product as an off-white solid. Purification by silica flash chromatography (50% EtOAc in hexanes) yielded NHS ester **2** (2.1874 g, 5.09 mmol, 84%) as a white solid.

**TLC**: $R_f$ 0.43 (50% EtOAc in hexanes). **$^1$H-NMR** (600 MHz): δ 5.22 (d, *J* = 8.7 Hz, 1H), 4.77 (s, 1H), 4.64 (q, *J* = 7.2 Hz, 1H), 3.14 (q, *J* = 6.6 Hz, 2H), 2.81 (s, 4H), 1.99 – 1.90 (m, 1H), 1.86 – 1.78 (m, 1H), 1.63 (br s, 2H), 1.40 (m, 18H). **$^{13}$C-NMR** (150 MHz): δ 168.8, 168.4, 156.1, 155.0, 80.6, 79.2, 51.8, 39.9, 29.9, 28.5, 28.3, 25.6. **HRMS** (ESI) *m/z* calcd for $C_{19}H_{31}N_3O_8Na$ ([M+Na]$^+$) 452.2009; found 452.2000.

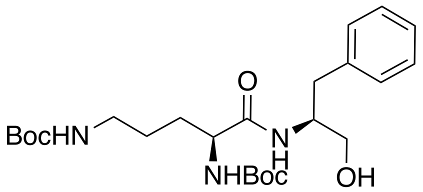


***N*[2],*N*[5]-Bis(*tert*-butoxycarbonyl)-L-ornithyl-L-phenylalaninol (3, Boc-Orn[Boc]-Phin-OH).** In a 25-mL roundbottom flask, L-phenylalaninol (148 mg, 0.978 mmol, 1.0 equiv) and DIPEA (405 µL, 2.94 mmol, 3.0 equiv) were dissolved in 10 mL anhyd THF and stirred at 0 °C for 10 min. *N*[2],*N*[5]-Bis(*tert*-butoxycarbonyl)-L-ornithine *N*-hydroxysuccinimide ester (**2**) (495 mg, 1.15 mmol, 1.18 equiv) was added to the stirring solution in one portion. The mixture was stirred at 0 °C for 5 min, warmed to 22 °C and stirred for 17 h. The mixture was diluted with $H_2O$ and extracted with $CH_2Cl_2$ (3 x 20 mL). The combined organic extracts were washed with brine, dried ($Na_2SO_4$), filtered, and concentrated by rotary evaporation to afford the crude product. Purification by silica flash chromatography (50% EtOAc in hexanes) yielded the amide **3** (378 mg, 0.81 mmol, 83%) as a white solid.

**TLC**: $R_f$ 0.25 (50% EtOAc in hexanes). **$^1$H-NMR** (600 MHz): δ 7.30 – 7.25 (m, 2H), 7.24 – 7.19 (m, 3H), 6.59 (d, *J* = 8.2 Hz, 1H), 5.24 (d, *J* = 7.6 Hz, 1H), 4.80 – 4.74 (m, 1H), 4.20 – 4.08 (m, 2H), 3.66 (dd, *J* = 11.3, 3.5 Hz, 1H), 3.54 (dd, *J* = 11.5, 4.8 Hz, 1H), 3.21 (s, 2H), 3.01 (dt, *J* = 13.8, 6.7 Hz, 1H), 2.87 (td, *J* = 12.2, 7.5 Hz, 2H), 1.80 – 1.71 (m, 1H), 1.62 – 1.46 (m, 2H), 1.42 (s, 18H). **$^{13}$C-NMR** (150 MHz): δ 172.3, 156.6, 155.9, 137.9, 129.4, 128.7, 126.7, 80.2, 79.6, 63.6, 54.0, 53.1, 39.8, 37.1, 30.1, 28.6, 28.4, 26.2. **HRMS** (ESI) *m/z* calcd for $C_{24}H_{39}N_3O_6Na$ ($[M+Na]^+$) 488.2737; found 488.2749.

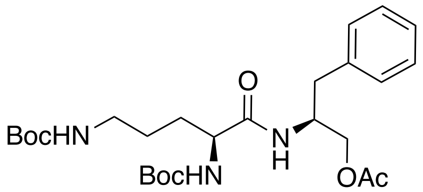


***N*[2],*N*[5]-Bis(tert-butoxycarbonyl)-L-ornithyl-L-phenylalaninol acetate (4, Boc-Orn[Boc]-Phin-OAc).** In a 10-mL roundbottom flask, *N*[2],*N*[5]-bis(*tert*-butoxycarbonyl)-L-ornithyl-L-phenylalaninol (**3**) (324 mg, 0.696 mmol, 1.0 equiv), 4-(dimethylamino)pyridine (18 mg, 0.15 mmol, 0.21 equiv), and triethylamine (194 µL, 1.39 mmol, 2.0 equiv) were dissolved in 5 mL anhyd $CH_2Cl_2$. Acetic anhydride (91 µL, 0.962 mmol, 1.38 equiv) was added to the stirring solution and the mixture was stirred at 22 °C for 21 h. The reaction was quenched by addition of satd aq $NaHCO_3$ and extracted with $CH_2Cl_2$ (3 x 20 mL). The combined organic extracts were washed with brine, dried ($Na_2SO_4$), filtered, and concentrated by rotary evaporation to afford the crude product. Purification by silica flash chromatography (50% EtOAc in hexanes) yielded the acetate **4** (325 mg, 0.64 mmol, 93%) as a white solid.

**TLC**: $R_f$ 0.50 (50% EtOAc in hexanes). **$^1$H-NMR** (600 MHz): δ 7.29 (t, *J* = 7.5 Hz, 2H), 7.24 – 7.17 (m, 3H), 6.50 (s, 1H), 5.08 (s, 1H), 4.67 (s, 1H), 4.40 (s, 1H), 4.12 (mf, 1H), 4.09 – 3.99 (m, 2H), 3.23 (s, 1H), 3.04 (dq, *J* = 12.5, 6.1 Hz, 1H), 2.84 (qd, *J* = 13.8, 7.1 Hz, 2H), 2.07 (s, 3H), 1.79 – 1.69 (m, 1H), 1.58 – 1.51 (m, 1H), 1.43 (m, 19H). **$^{13}$C-NMR** (150 MHz): δ 171.8, 171.3, 171.1, 156.5, 155.8, 137.1, 129.3, 128.8, 126.9, 80.1, 79.4, 64.8, 60.5, 53.6, 53.6, 49.7, 39.6, 37.6, 30.1, 28.6, 28.4, 26.4, 20.9, 14.3. **HRMS** (ESI) *m/z* calcd for $C_{26}H_{42}N_3O_7$ ($[M+H]^+$) 508.3023; found 508.3023.

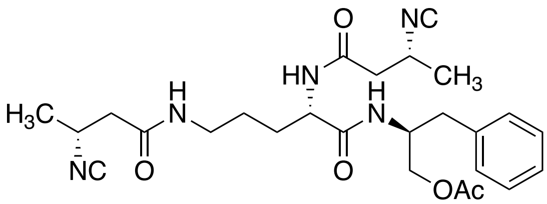


**$N^2$,$N^5$-Bis((*R*)-3-isocyanobutanoyl)-L-ornithyl-L-phenylalaninol acetate (6, C4-Orn-Phin-OAc diisonitrile).** In a 20-mL glass vial, $N^2$,$N^5$-bis(*tert*-butoxycarbonyl)-L-ornithyl-L-phenylalaninol acetate (**4**) (350 mg, 0.689 mmol, 1.0 equiv) was dissolved in 8 mL anhyd $CH_2Cl_2$ and stirred at 0 °C. Trifluoroacetic acid (2 mL, 25.6 mmol, 37.1 equiv) was added and the mixture was stirred at 0 °C for 5 min, warmed to 22 °C and stirred for 1 h. The mixture was concentrated by rotary evaporation to afford the crude bis(trifluoroacetate) salt (**5**).

In a 20-mL roundbottom flask, the crude bis(trifluoroacetate) salt (**5**) (369 mg, 0.689 mmol, 1.0 equiv) and DIPEA (1.08 mL, 6.20 mmol, 9 equiv) were dissolved in 5 mL anhyd $CH_2Cl_2$ and stirred at 0 °C. 2,5-Dioxopyrrolidin-1-yl (*R*)-3-isocyanobutanoate, prepared as previously described,[2] (367 mg, 1.75 mmol, 2.53 equiv) was added to the stirring solution and the mixture was stirred at 0 °C for 5 min, warmed to 22 °C and stirred for 17 h. The mixture was diluted with $H_2O$ and extracted with $CH_2Cl_2$ (3x20 mL). The combined organic extracts were washed with brine, dried ($Na_2SO_4$), filtered, and concentrated by rotary evaporation to afford the crude product. Purification by silica flash chromatography (100% EtOAc) yielded the diisonitrile **6** (212 mg, 0.43 mmol, 62%) as a colorless opaque solid.

**TLC**: $R_f$ 0.18 (100% EtOAc). **$^1$H-NMR** (600 MHz, DMSO-$d_6$): δ 8.15 (d, $J$ = 8.2 Hz, 1H), 8.02 (t, $J$ = 5.7 Hz, 1H), 7.99 (d, $J$ = 8.4 Hz, 1H), 7.27 (m, 2H), 7.20 (m, 3H), 4.25 (q, $J$ = 7.6 Hz, 1H), 4.16 (q, $J$ = 6.8 Hz, 1H), 4.06 (m, 2H), 4.01 (dd, $J$ = 11.1, 4.7 Hz, 1H), 3.84 (dd, $J$ = 11.1, 7.1 Hz, 1H), 3.03 (dh, $J$ = 20.0, 6.7 Hz, 2H), 2.77 (dd, $J$ = 13.9, 6.2 Hz, 1H), 2.72 (dd, $J$ = 13.8, 7.8 Hz, 1H), 2.56 (dd, $J$ = 14.6, 8.2 Hz, 1H), 2.49 – 2.35 (m, 3H), 1.99 (s, 3H), 1.56 (m, 1H), 1.50 – 1.33 (m, 1H), 1.30 (t, $J$ = 7.1 Hz, 6H). **$^{13}$C-NMR** (150 MHz, DMSO-$d_6$): δ 171.1, 170.3, 168.0, 167.9, 155.0, 154.9, 154.9, 137.9, 129.1, 128.2, 126.2, 64.6, 52.2, 48.9, 46.99, 46.95, 46.92, 42.2, 42.0, 38.2, 36.6, 29.9, 25.5, 20.99, 20.95, 20.6. **HRMS** (ESI) *m/z* calcd for $C_{26}H_{35}N_5O_5Na$ ($[M+Na]^+$) 520.2536; found 520.2536.

[2] Xu, Y.; Tan, D. S. *Org. Lett.* **2019**, *21*, 8731-8735.

#### B. $^{1}$H-NMR AND $^{13}$C-NMR SPECTRA

**1. SYNTHESIS OF DIISONITRILE 6** **S7**

a. Boc-Orn(Boc)-NHS (**2**) S7
b. Boc-Orn(Boc)-Phin-OH (**3**) S8
c. Boc-Orn(Boc)-Phin-OAc (**4**) S9
d. C4-Orn-Phin-OAc Diisonitrile (**6**) S10

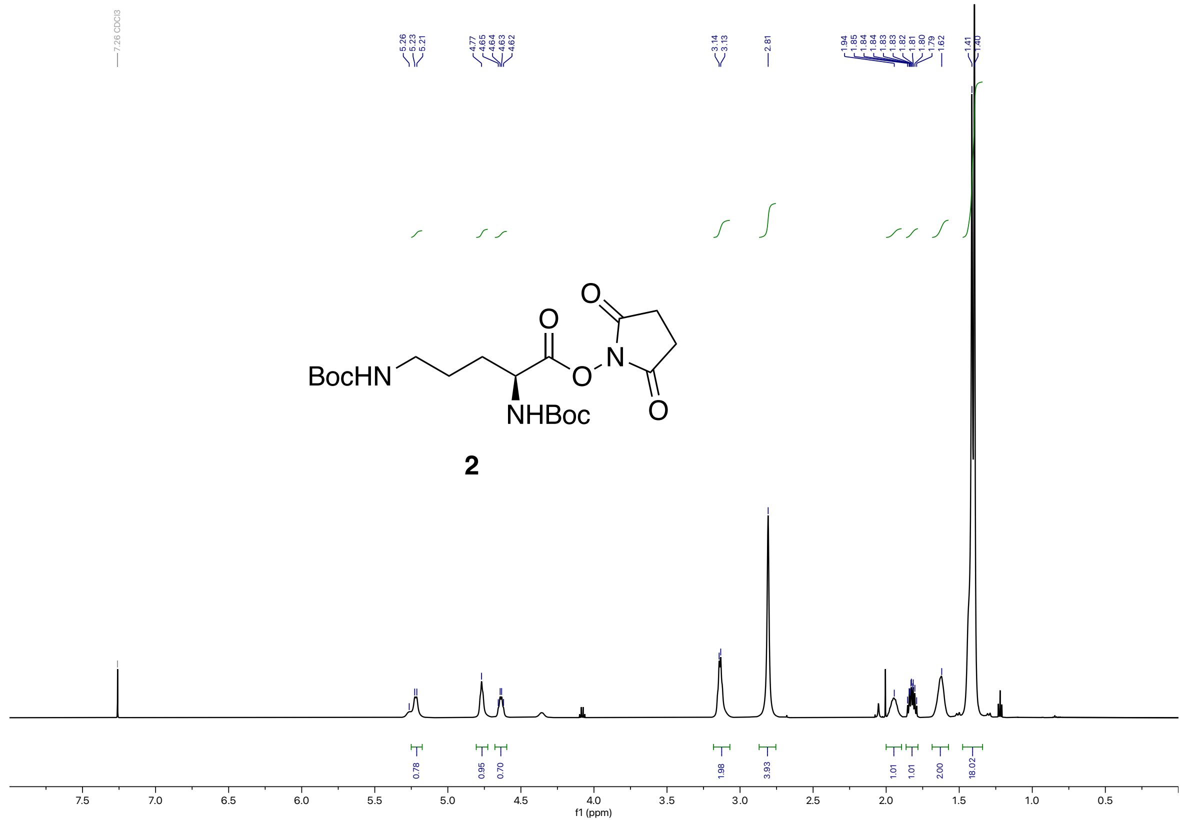
BocHN
NHBoc
2
f1 (ppm)

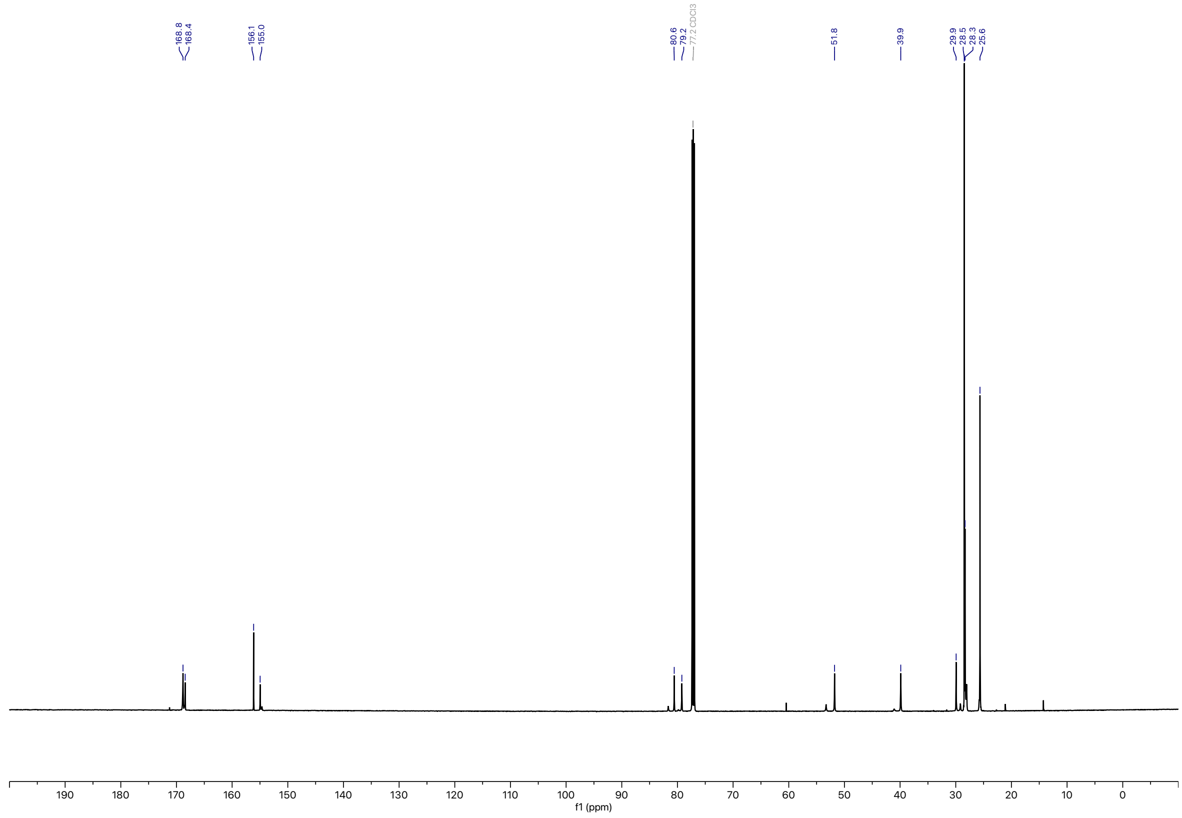
168.8
168.4
156.1
155.0
80.6
79.2
77.2 CDCl3
51.8
39.9
29.9
28.5
28.3
25.6
f1 (ppm)

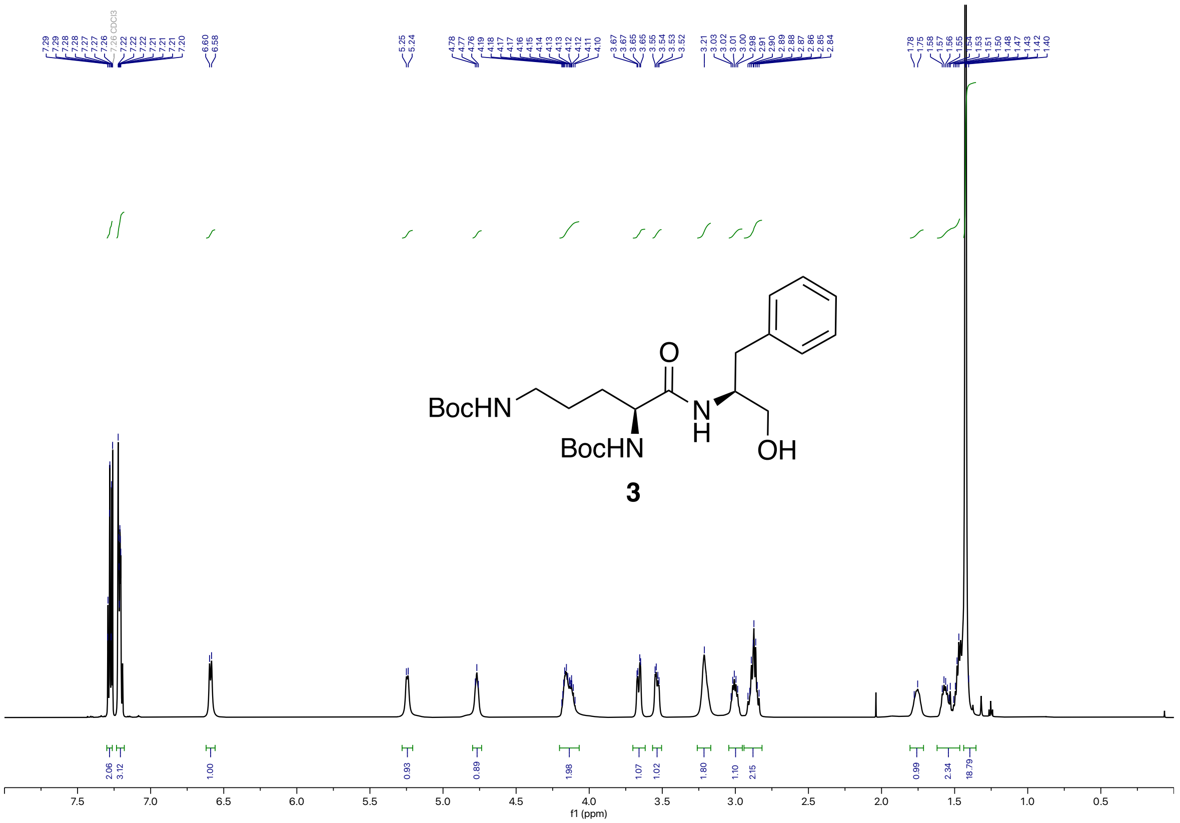
BocHN
BocHN
O
N
H
OH
3
f1 (ppm)

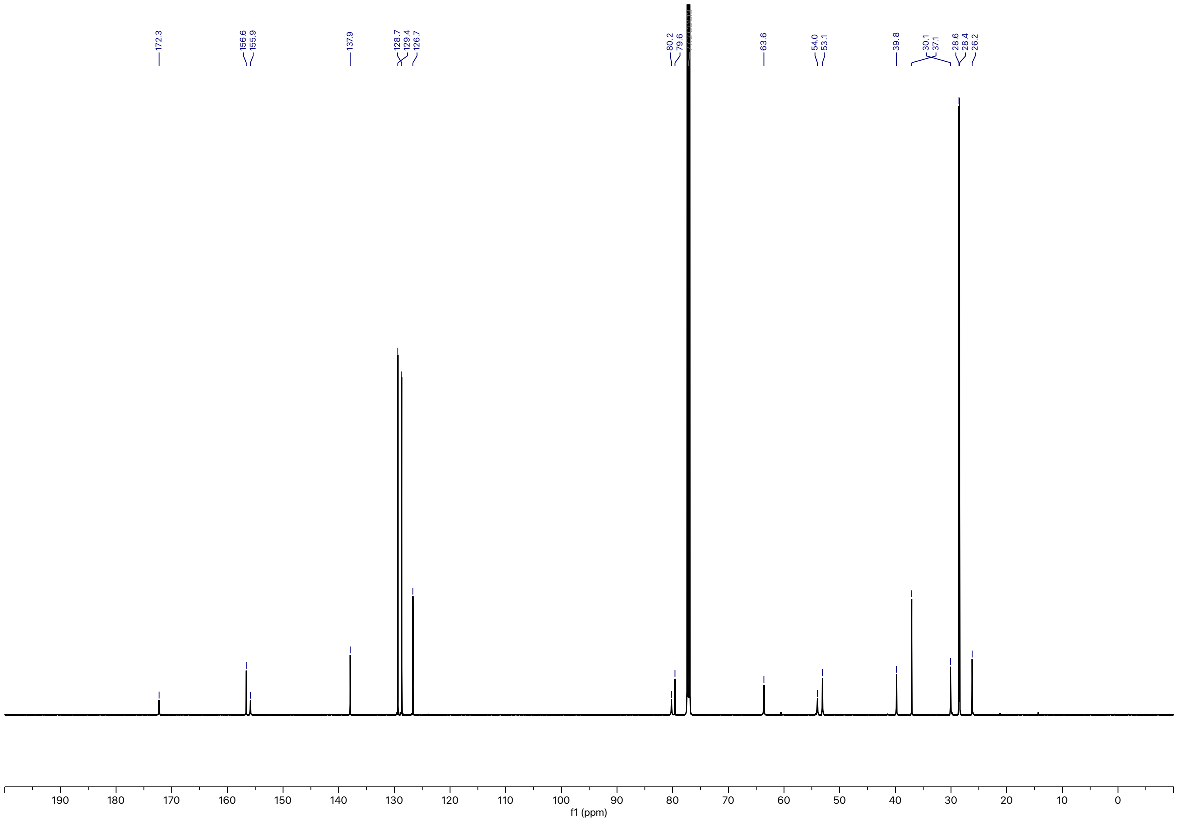
f1 (ppm)

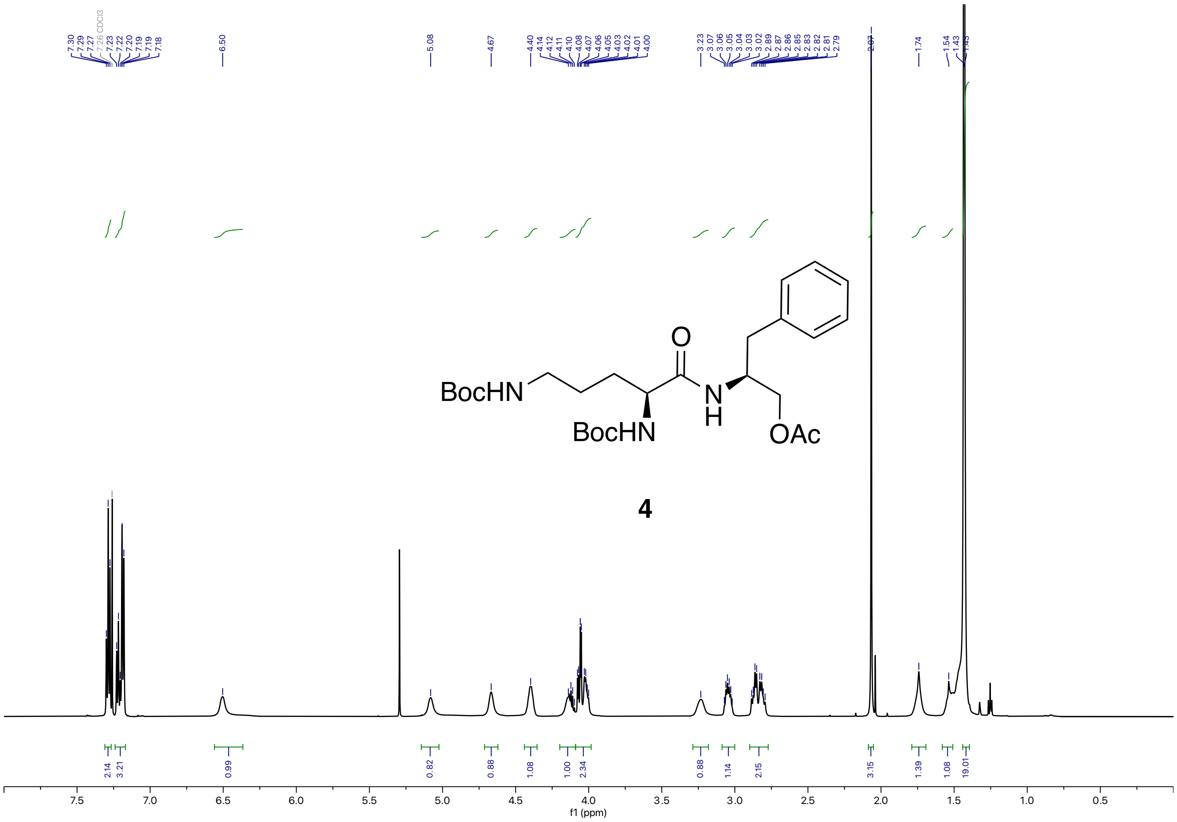
7.30
7.29
7.27
7.26 CDCl3
7.23
7.22
7.20
7.19
7.18
6.50
5.08
4.67
4.40
4.14
4.12
4.11
4.10
4.08
4.07
4.06
4.05
4.03
4.02
4.01
4.00
3.23
3.07
3.06
3.05
3.04
3.03
3.02
2.89
2.87
2.86
2.85
2.83
2.82
2.81
2.79
2.07
1.74
1.54
1.43
O
BocHN
N
H
BocHN
OAc
4
2.14
3.21
0.99
0.82
0.88
1.08
1.00
2.34
0.88
1.14
2.15
3.15
1.39
1.08
19.01
7.5
7.0
6.5
6.0
5.5
5.0
4.5
4.0
3.5
3.0
2.5
2.0
1.5
1.0
0.5
f1 (ppm)

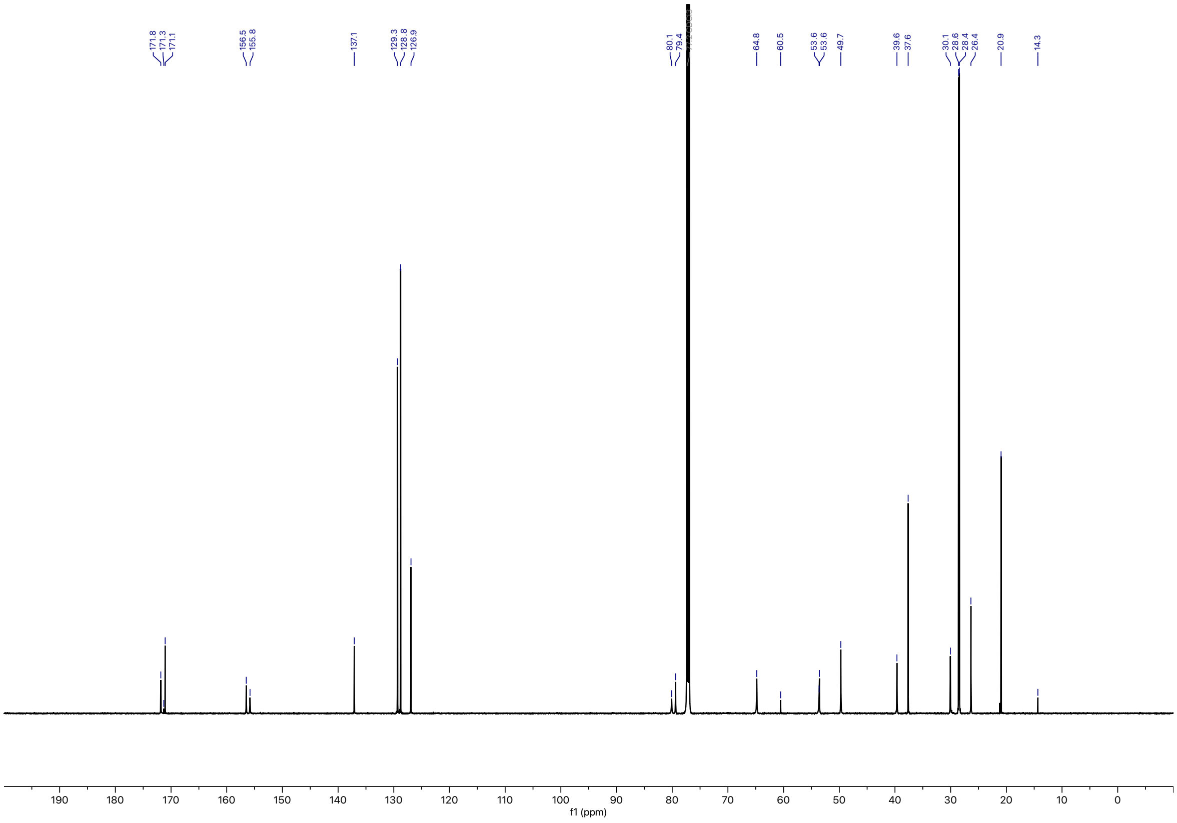
171.8
171.3
171.1
156.5
155.8
137.1
129.3
128.8
126.9
80.1
79.4
64.8
60.5
53.6
53.6
49.7
39.6
37.6
30.1
28.6
28.4
26.4
20.9
14.3
190
180
170
160
150
140
130
120
110
100
90
80
70
60
50
40
30
20
10
0
f1 (ppm)

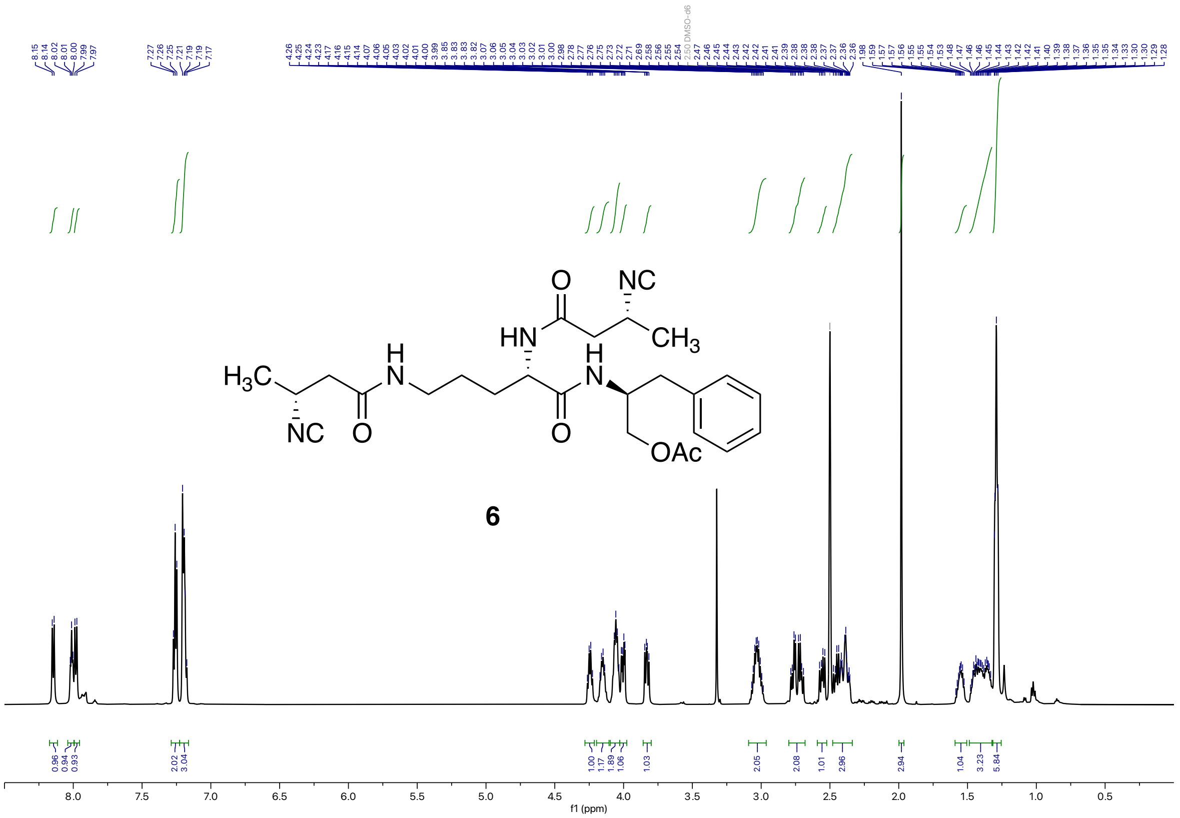

**6**

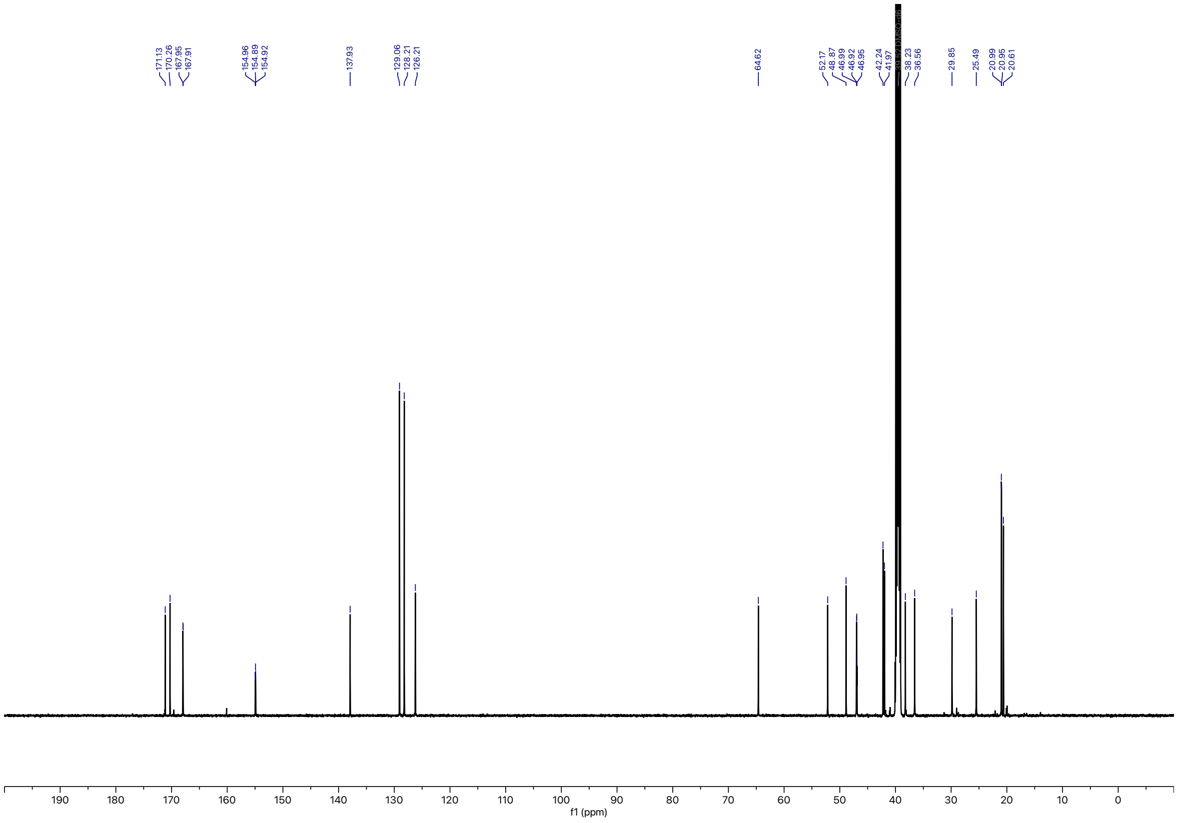
